# Cerebrovascular damage caused by the gut microbe/host co-metabolite *p*-cresol sulfate is prevented by blockade of the EGF receptor

**DOI:** 10.1101/2022.11.12.516113

**Authors:** Sita N. Shah, Tobias B-A. Knausenberger, Matthew G. Pontifex, Emily Connell, Gwénaëlle Le Gall, Tom A. J. Hardy, David W. Randall, Kieran McCafferty, Muhammad M. Yaqoob, Egle Solito, Michael Müller, Andrew V. Stachulski, Robert C. Glen, David Vauzour, Lesley Hoyles, Simon McArthur

**Affiliations:** Blizard Institute, Faculty of Medicine & Dentistry, Queen Mary, University of London, 4, Newark Street, London, UK; Norwich Medical School, Biomedical Research Centre, University of East Anglia, Norwich, UK; Department of Renal Medicine and Transplantation, Royal London Hospital, Barts Health NHS Trust, London, UK; William Harvey Research Institute, Faculty of Medicine & Dentistry, Queen Mary, University of London, John Vane Science Centre, Charterhouse Square, London, UK; Dipartimento di Medicina Molecolare e Biotecnologie Mediche, Università degli Studi di Napoli Federico II, Naples, Italy; Robert Robinson Laboratories, Department of Chemistry, University of Liverpool, Liverpool, UK; Faculty of Medicine, Department of Metabolism, Digestion and Reproduction, Imperial College London, London, UK; Centre for Molecular Informatics, Department of Chemistry, University of Cambridge, Cambridge, UK; Department of Biosciences, School of Science and Technology, Nottingham Trent University, Clifton, Nottingham, UK; Institute of Dentistry, Faculty of Medicine & Dentistry, Queen Mary, University of London, Blizard Institute, 4, Newark Street, London, UK

**Keywords:** Blood-brain barrier, *p*-cresol sulfate, cerebrovascular disease, chronic kidney disease, gut microbiota

## Abstract

Circulating levels of the gut microbe/host co-metabolite p-cresol sulfate (pCS) correlate with cerebrovascular event risk in individuals with chronic kidney disease, but whether this relationship is mechanistic is unclear. We hypothesised that pCS would impair function of the blood-brain barrier (BBB), the primary brain-vasculature interface. We report that pCS exposure impairs BBB integrity in human cells *in vitro* and both acutely and chronically in mice, enhancing tracer extravasation, disrupting barrier-regulating tight junction components and ultimately affecting whole-brain transcriptomic activity. *In vitro* and *in vivo* mechanistic studies showed that pCS activated epidermal growth factor receptor (EGFR) signalling, resulting in matrix metalloproteinase mobilisation and BBB damage. Furthermore, blockade of EGFR prevented the permeabilising effects of serum from haemodialysis patients upon cerebromicrovascular endothelia *in vitro*. Our results define a pathway linking the co-metabolite pCS with BBB damage and suggest targeting the EGFR may mitigate against cerebrovascular damage in CKD.

---

Communication between the commensal microbiota and host systems is implicated in many areas of health and disease^1^, with substantial evidence highlighting gut microbes as modifiers of CNS function^2^, but far less known about how these links are mediated^3^. Microbial modulation of afferent neural^4^, immune^5^ and neuroendocrine^6^ activity has been described, but perhaps the least well characterised route of gut-brain communication is direct exposure to the myriad microbe-derived metabolites present in the circulation^7^. We and others have recently shown gut microbe-derived circulating metabolites (e.g. short-chain fatty acids^8,9^ and methylamines^10^) to target the blood-brain barrier (BBB), but the impact of other gut microbial metabolites upon the brain vasculature, and their roles in health and disease, are largely unexplored.

One important microbe-derived metabolite given its links with kidney dysfunction is *p*-cresol, produced by fermentation of tyrosine and phenylalanine in the colon, particularly by *Coriobacteriaceae* or *Clostridium* species^11,12^. Luminal *p*-cresol undergoes extensive conjugation in both enterocytes^13^ and the liver^14^, such that it is found almost exclusively as *p*- cresol sulfate (pCS, ∼90 % in humans) and *p*-cresol glucuronide (pCG) in the systemic circulation^15^. Normally, these molecules are efficiently cleared by the kidneys with an effective plasma half-life of ∼30 min in rodents^16^. However, due to its high binding affinity to albumin, pCS accumulates within the plasma of individuals with compromised renal function^17^ and is poorly removed by haemodialysis^18^, a feature of significance given that increased plasma concentrations of pCS predict mortality in individuals with chronic kidney disease (CKD)^19,20^.

CKD is a complex disorder affecting many aspects of physiology, but is notably associated with significant neurological and cerebrovascular complications^21^. Patients with CKD are at an increased risk of both ischaemic and haemorrhagic stroke, even after correcting for coincident vascular risk factors^22^, and tend to have worse functional outcomes post-infarct^23^, a major unmet clinical need^24^. Moreover, CKD is associated with both enhanced BBB permeability^25^ and the presence of microbleeds^26^, suggesting the cerebral vasculature is directly damaged in the condition.

These links between CKD and cerebrovascular disease, alongside reports from clinical studies associating plasma pCS and stroke risk^27^ and experimental work showing pCS to induce dermal microvascular leakage^28^ and aortic oxidative damage and remodelling^28,29^, led us to hypothesise that this gut microbe-host co-metabolite directly damages the cerebral vasculature, impairing BBB function and contributing to the neurological consequences of CKD. Here, we report a clear deleterious effect of pCS upon BBB integrity, mediated through stimulation of the epidermal growth factor receptor (EGFR) and ensuing matrix metalloproteinases (MMPs) activation. Moreover, we show antagonism of the EGFR to prevent the BBB-permeabilising actions of serum from CKD patients, offering a potential new pathway to protect against uraemia-associated cerebrovascular damage linked to the actions of the gut microbiota.

## Results

### Treatment with pCS increases brain endothelial permeability in vitro and in vivo

To assess the effects of pCS upon BBB integrity, we first examined its effects upon a simple *in vitro* model of BBB function, namely the paracellular permeability barrier of a monolayer of the immortalised human cerebromicrovascular endothelial line hCMEC/D3^30^. Treatment with pCS (24 h; 10 µM, 100 µM or 1 mM) significantly (*P* < 0.05) and dose-dependently enhanced permeability to a 70 kDa FITC-dextran tracer (Fig. 1A) and reduced trans-endothelial electrical resistance (TEER; Fig. 1B). Notably, pCS showed no signs of endothelial toxicity at any concentration tested (Extended Data Fig. 1) and given that pCS was administered as a potassium salt, equivalent concentrations of KCl had no effect on barrier function (Extended Data Fig. 1). Moreover, chelation of pCS by inclusion of 50 mg/ml human albumin (approximately equivalent to human serum concentrations^31^) blocked its effects on paracellular permeability and TEER (Extended Data Fig. 2). The BBB permeability barrier is underpinned by tight inter-endothelial junctions and their interactions with the actin cytoskeleton; hence, we examined the effect of pCS treatment upon the key tight junction component zonula occludens-1 (ZO-1) and fibrillar actin in hCMEC/D3 monolayers. In comparison with untreated cells, pCS exposure (24 h, 10 µM) markedly disrupted the marginal localisation of ZO-1 (Fig. 1C), as well as disrupting actin fibril arrangement itself (Fig. 1D), with a clear loss of cortical actin and appearance of trans-cytoplasmic fibres.

**Figure 1:**
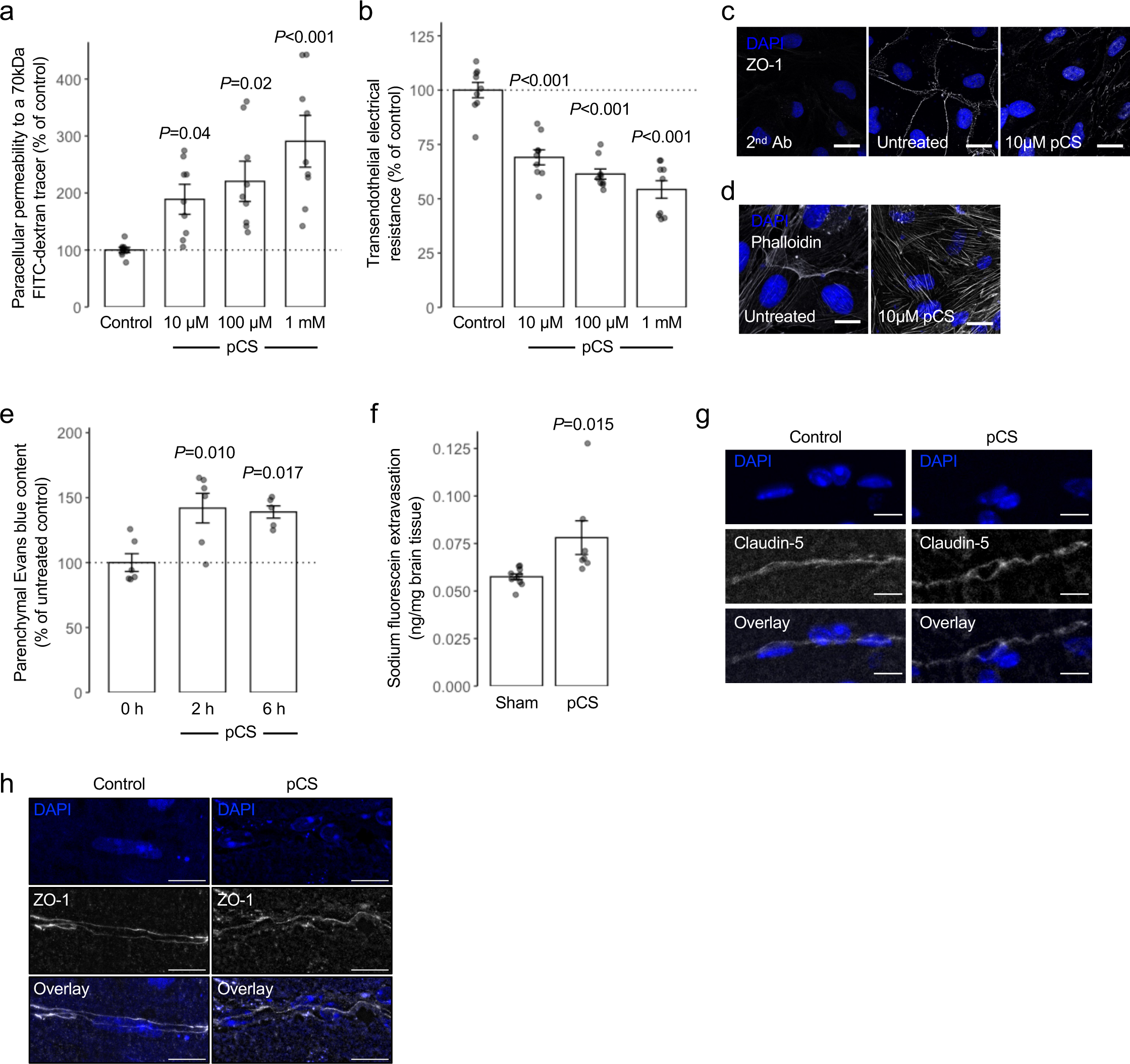
Exposure to pCS increases permeability of human cerebromicrovascular endothelial cells *in vitro* and the murine BBB *in vivo*. a) Incubation of hCMEC/D3 cell monolayers with pCS (10 µM, 100 µM, 1 mM; 24 h) dose-dependently increased paracellular permeability to a 70 kDa FITC-dextran conjugate; data are mean ± s.e.m., n = 9 independent experiments. b) Incubation of hCMEC/D3 cell monolayers with pCS (10 µM, 100 µM, 1 mM; 24 h) dose-dependently reduced TEER; data are mean ± s.e.m., n = 9 independent experiments. c, d) Confocal microscopic analysis of expression of c) AF488-phalloidin labelled actin filaments (white) or d) the tight junction component zona occludens-1 (ZO-1; white) in hCMEC/D3 cells following treatment for 24 h with 10 µM pCS, nuclei are counterstained with DAPI (blue), scale = 15 µm. Images are representative of at least three independent experiments. e) Treatment of male C57Bl/6 mice by i.p. injection of pCS (10 mg/kg) caused a time-dependent increase in extravasation of Evans blue tracer into the CNS parenchyma, reaching statistical significance at both 2 h and 6 h post administration; data are mean ± s.e.m., n = 5-6 animals. f) Exposure of male C57Bl/6 by s.c. implantation with osmotic mini-pumps releasing pCS at 7.5 µg/h for 28 days significantly enhanced extravasation of sodium fluorescein into the brain parenchyma; data are mean ± s.e.m., n = 7-10 animals. g, h) Confocal microscopic analysis of expression of g) claudin-5 or h) zonula occludens-1 (ZO-1) in the brains of male C57Bl/6 exposed by s.c. implantation with osmotic mini-pumps releasing pCS at 7.5 µg/h for 28 days, nuclei are counterstained with DAPI (blue), scale = 10 µm. Images are representative of 7-10 animals.

To confirm that these *in vitro* findings had relevance for whole animal physiology, we exposed male C57Bl/6 mice to pCS (10 mg/kg i.p., chosen to approximately double circulating free pCS concentrations^32^; 2 h, 6 h) before assessing BBB permeability through cerebral extravasation of the albumin-binding tracer Evans blue. Treatment of mice with pCS resulted in a significant (*P* < 0.02) increase in brain parenchymal Evans blue at both 2 h and 6 h post-administration (Fig. 1E).

CKD is characterised in part by persistently elevated circulating levels of pCS; hence we examined how continuous pCS release from implanted osmotic minipumps for four weeks affected the BBB, neuroinflammation and cognitive function (see Extended Data Table 1 for serum metabolite content analysis). Animals so exposed to pCS exhibited both significantly enhanced extravasation of the tracer sodium fluorescein into the brain parenchyma (Fig. 1F), and disrupted expression of the key BBB inter-endothelial tight junction molecules claudin-5 (Fig. 1G) and ZO-1 (Fig. 1H), confirming the deleterious effects seen with acute pCS administration. However, no obvious sign of microglial or astrocytic change were seen in the cortex of these animals (Extended Data Fig. 3), nor was performance in the open field test, Y-maze or novel object recognition task significantly affected (Extended Data Fig. 4) in comparison to saline-treated animals.

**Table 1:** Significant results generated from SPIA of 380 significantly differentially expressed genes.

| **Name** | **KEGG**<br>**pathway ID** | **pGFdr** | **pGFWER** | **Status** |
| --- | --- | --- | --- | --- |
| Glutamatergic synapse | mmu:04724 | $2.83 \times 10^{-4}$ | $2.83 \times 10^{-4}$ | Inhibited |
| Circadian entrainment | mmu:04713 | $2.98 \times 10^{-3}$ | $5.97 \times 10^{-3}$ | Inhibited |
| Alzheimer disease | mmu:05010 | $1.40 \times 10^{-2}$ | $4.20 \times 10^{-2}$ | Inhibited |
| Amphetamine addiction | mmu:05031 | $1.92 \times 10^{-2}$ | $7.68 \times 10^{-2}$ | Inhibited |
| Oxytocin signalling pathway | mmu:04921 | $2.54 \times 10^{-2}$ | $1.27 \times 10^{-1}$ | Inhibited |
| Glucagon Signaling pathway | mmu:04922 | $2.75 \times 10^{-2}$ | $1.72 \times 10^{-1}$ | Inhibited |
| Retrograde endocannabinoid<br>signalling | mmu:04723 | $2.75 \times 10^{-2}$ | $1.92 \times 10^{-1}$ | Inhibited |
| Pathways of neurodegeneration -<br>multiple diseases | mmu:05022 | $2.98 \times 10^{-2}$ | $2.66 \times 10^{-1}$ | Inhibited |
| VEGF signalling pathway | mmu:04370 | $2.98 \times 10^{-2}$ | $2.68 \times 10^{-1}$ | Inhibited |
| Cellular senescence | mmu:04218 | $3.12 \times 10^{-2}$ | $3.13 \times 10^{-1}$ | Inhibited |
| Aldosterone synthesis and secretion | mmu:04925 | $3.12 \times 10^{-2}$ | $3.43 \times 10^{-1}$ | Inhibited |
| Ras signalling pathway | mmu:04014 | $4.14 \times 10^{-2}$ | $4.96 \times 10^{-1}$ | Inhibited |
| C-type lectin receptor signalling<br>pathway | mmu:04625 | $4.18 \times 10^{-2}$ | $5.44 \times 10^{-1}$ | Inhibited |
| Coronavirus disease - COVID-19 | mmu:05171 | $4.59 \times 10^{-2}$ | $6.42 \times 10^{-1}$ | Activated |
| Lipid and atherosclerosis | mmu:05417 | $4.83 \times 10^{-2}$ | $7.24 \times 10^{-1}$ | Inhibited |
| Gastric acid secretion | mmu:04971 | $4.94 \times 10^{-2}$ | $7.90 \times 10^{-1}$ | Inhibited |
| Huntington disease | mmu:05016 | $4.97 \times 10^{-2}$ | $8.45 \times 10^{-1}$ | Inhibited |

### Murine whole brain transcriptomic analysis reveals a suppressive effect of pCS exposure upon neuronal activity

To investigate potential mechanisms of pCS action, we performed RNAseq analysis of whole-brain transcriptomes, comparing untreated and pCS-exposed mice (10 mg/kg i.p., 2 h). We found 380/16,988 genes assayed were significantly differentially expressed (*P* < 0.05, Benjamini-Hochberg): with 91 genes being significantly upregulated in the pCS-exposed group, while 289 were significantly downregulated (Fig. 2A; Extended Data Table 2). Among BBB-specific genes, the expression of *Slc1a2* (log2 fold change -1.35), *Adgra2* (1.03) and *Serpini1* (-1.93) was found to be significantly changed (*P* <2.66x10^-4^ for all, Benjamini- Hochberg).

**Figure 2:**
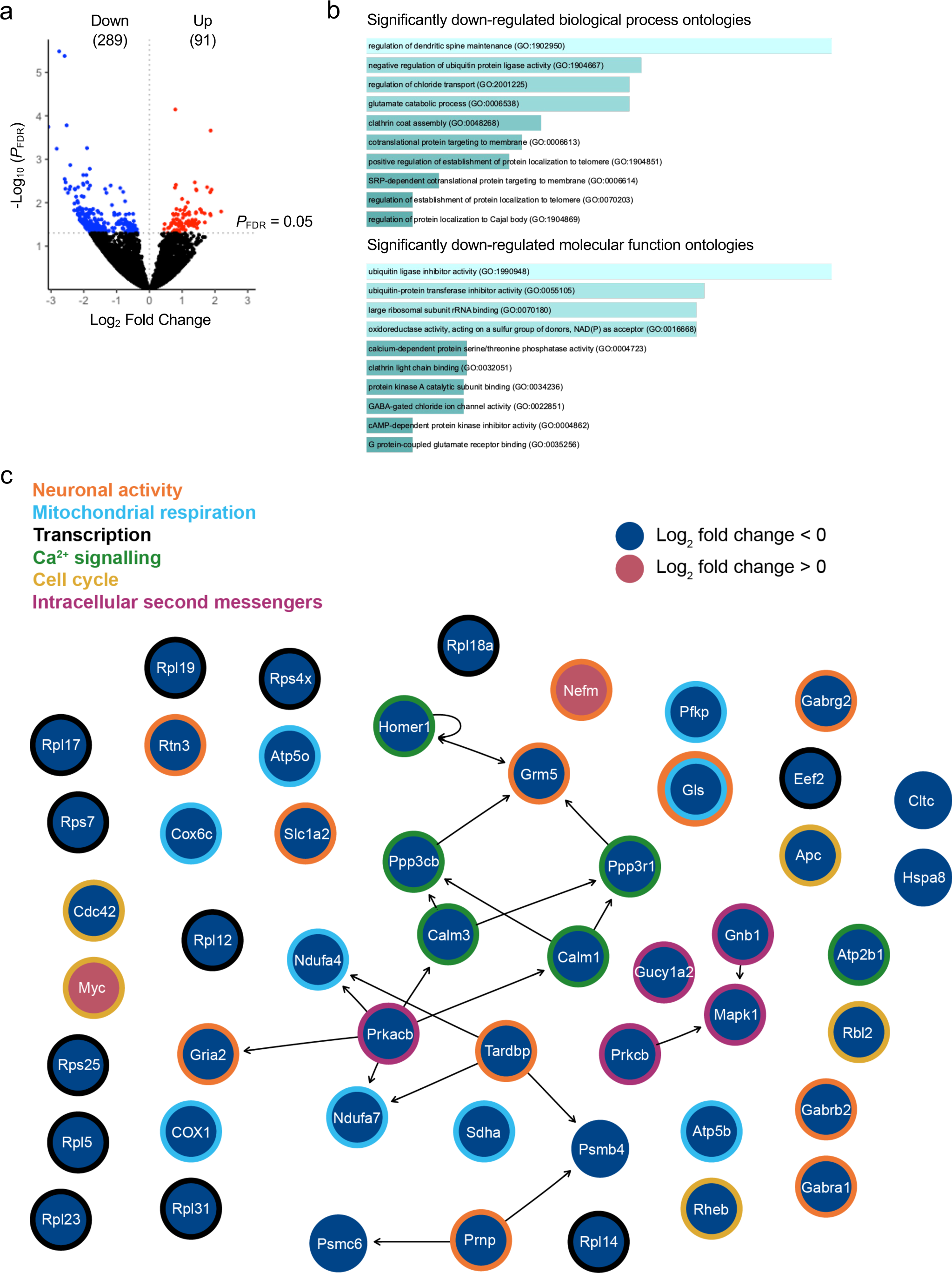
Exposure of mice to pCS for 2 h significantly affects brain gene expression. a) Volcano plot showing the 380 significantly differentially expressed genes among the 16,988 mouse genes examined in this study (n = 5 animals per group (treated, negative control)). Black data points: no significant change in expression; red dots, significantly (*P* < 0.05, Benjamini-Hochberg) upregulated expression in the pCS group compared with the negative control; blue dots, significantly (*P* < 0.05, Benjamini-Hochberg) downregulated expression in the pCS group compared with the negative control; n=5 per group. b) Biological processes (above) and molecular functions (below) of genes found to be significantly downregulated (n= 289) upon exposure of mice to pCS, based on Enrichr *P* value ranking from Gene Ontology analysis. c) Topological analysis of the KEGG network resulting from mapping the 380 significantly (*P* < 0.05, Benjamini-Hochberg) differentially expressed genes onto the 17 KEGG *Mus musculus* metabolic pathways listed in Table 1. Blue solid circles, genes whose expression is significantly downregulated in the pCS group compared with the control; red solid circles, genes whose expression is significantly upregulated in the pCS group compared with the control. Thick outline colours around the solid circles correspond to the molecular functions listed in the legend. Network summary statistics are available in Table S3.

**Table 2:** Serum donor demographics.

|  | Age<br>(years) | Sex<br>ratio | Serum urea<br>(mg/dl) | Serum<br>creatinine (μM) | Renal replacement<br>therapy duration<br>(years) |
| --- | --- | --- | --- | --- | --- |
| Healthy donor | 49 ± 3.3 | 8F : 3M | 4.46 ± 0.36 | 82.0 ± 6.8 | N/A |
| Haemodialysis | 59 ± 6.2 | 4F : 6M | 19.13 ± 1.90 | 876.2 ± 94.7 | 10.6 ± 4.2 |

Analysis of over-represented Gene Ontology categories within differentially expressed genes using Enrichr^33^ revealed that, while no categories associated with significantly up-regulated genes reached statistical significance after multiple testing correction, several biological processes and molecular functions associated with significantly (*P* < 0.05; Benjamini-Hochberg) down-regulated genes could be identified (Fig. 2B). Broadly, these fell into two main categories: ontologies reflective of neuronal function and synaptic activity, and those indicating intracellular protein translocation and metabolism. Signalling Pathway Impact Analysis (SPIA) demonstrated inhibition of pathways associated with glutamatergic synapse, circadian entrainment, and Alzheimer disease (pGFWER < 0.05; Table 1). Topological analysis of the network generated from the 380 significantly differentially expressed genes mapped onto KEGG pathways identified in SPIA (pGFdr < 0.05; Table 1) suggested that the genes for *Prkacb*, *Homer1*, *Tardbp*, *Calm1*, *Calm3*, *Ppp3r1*, *Ppp3cb* and *Grm5* had most control over the network (Fig. 2C; Extended Data Table 3). Manual curation of SPIA- highlighted genes identified several broad categories of primarily down-regulated function: neuronal activity, mitochondrial respiration, translation, cell cycle activity, and intracellular Ca^2+^- and other signalling pathways (Fig. 2C), overall depicting a generalised inhibitory effect of pCS upon brain function.

**Table 3:** Human haemodialysis patient serum metabolite concentrations. LC-MS/MS determined concentrations of different potential uraemic toxins in sera from healthy donors and individuals undergoing haemodialysis for nephropathy, alongside coefficients of correlation between metabolite concentration and permeability of hCMEC/D3 cells to a 70 kDa FITC-dextran tracer following treatment with 20% sera for 24 h; n = 10-11, * *P*<0.05 *vs.* healthy donors.

| Metabolite | Healthy donor<br>($\mu\text{M}$) | Haemodialysis<br>($\mu\text{M}$) | Permeability<br>correlation | |
| --- | --- | --- | --- | --- |
| | | | Rho | $P_{\text{FDR}}$ |
| *p*-Cresol sulfate | 14.12 ± 3.09 | 155.26 ± 27.36 * | 0.675 | 0.01 |
| *p*-Cresol glucuronide | 0.09 ± 0.03 | 10.63 ± 4.44 * | 0.394 | 0.12 |
| Indoxyl sulfate | 2.92 ± 0.45 | 147.18 ± 26.46 * | 0.400 | 0.09 |
| Serotonin | 0.64 ± 0.12 | 0.42 ± 0.18 | -0.233 | 0.64 |
| Kynurenine | 1.73 ± 0.23 | 4.63 ± 0.57 * | 0.371 | 0.06 |
| 3-Hydroxyanthranilic acid | 0.01 ± 0.00 | 0.05 ± 0.01 * | 0.434 | 0.11 |
| Tryptophan | 42.30 ± 1.46 | 22.97 ± 2.35 * | -0.466 | 0.06 |
| 5-Hydroxyindole acetic acid | 0.05 ± 0.01 | 1.28 ± 0.21 * | 0.464 | 0.08 |
| Xanthurenic acid | 0.01 ± 0.00 | 0.17 ± 0.05 * | 0.442 | 0.02 |
| Anthranilic acid | 0.20 ± 0.02 | 0.52 ± 0.1 * | 0.396 | 0.12 |
| Kynurenic acid | 0.06 ± 0.01 | 2.34 ± 0.58 * | 0.353 | 0.16 |
| Indole-3-lactic acid | 0.78 ± 0.12 | 2.42 ± 0.33 * | 0.397 | 0.12 |
| Indole-3-carboxaldehyde | 0.05 ± 0.00 | 0.06 ± 0.01 | -0.08 | 0.78 |
| Indole-carboxylic acid | 0.04 ± 0.01 | 0.06 ± 0.01 | 0.041 | 0.89 |
| Indole-3-acetic acid | 2.45 ± 0.75 | 6.36 ± 2.07 | 0.385 | 0.17 |
| Indole-3-propionic acid | 1.73 ± 0.4 | 1.37 ± 0.35 | 0.051 | 0.47 |
| Methyl indole 3-acetate | 0.05 ± 0.03 | 7.44 ± 3.48 * | 0.241 | 0.43 |

### Permeabilising effects of pCS are mediated through the EGFR

Although transcriptomic analysis did not suggest a clear candidate pathway for the actions of pCS directly upon the cerebral vasculature, previous studies have indicated that it may interact with the EGFR^34^, hence we investigated the potential involvement of this receptor in the effects of the metabolite upon the BBB. Initial studies of hCMEC/D3 cells confirmed EGFR expression (Fig. 3A) and showed it to be phosphorylated at Tyr1068 within 15 min of 10 µM pCS stimulation, phosphorylation that was maintained for at least 30 min (Fig. 3B). That this was a direct effect of pCS was confirmed by inclusion of the EGFR-specific inhibitor erlotinib (2.5 µM, 10 min pre-treatment), which effectively prevented EGFR phosphorylation (Fig. 3C).

**Figure 3:**
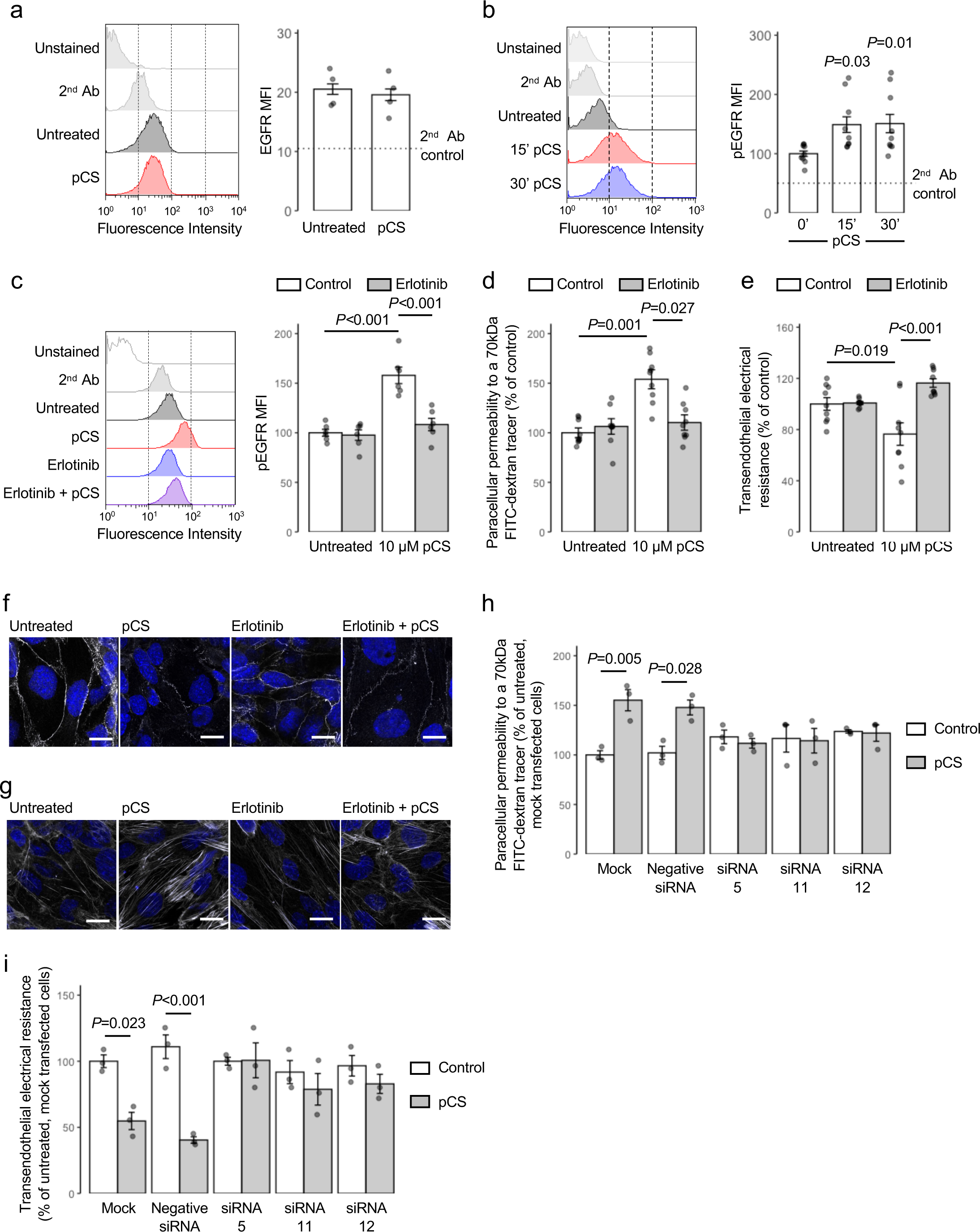
pCS regulates cerebromicrovascular cell permeability *in vitro* through activation of the EGFR. a) Stimulation of hCMEC/D3 cells with pCS (24 h, 10 µM) has no effect on cell surface expression of EGFR; data are mean ± s.e.m., n = 4 independent experiments; representative flow cytometry histograms are shown. b) Treatment of hCMEC/D3 cells with pCS (10 µM) causes a time-dependent increase in EGFR Tyr1068 phosphorylation, maintained for at least 30 min; data are mean ± s.e.m., n = 9 independent experiments; representative flow cytometry histograms are shown. c) Pre-treatment of hCMEC/D3 cells with erlotinib (2.5 µM, 10 min) prevents EGFR Tyr1068 phosphorylation in response to pCS treatment (10 µM, 30 min), data are mean ± s.e.m., n = 6 independent experiments; representative flow cytometry histograms are shown. d) Pre-treatment of hCMEC/D3 cell monolayers with erlotinib (2.5 µM, 10 min) prevents the increase in paracellular permeability to a 70 kDa FITC-dextran conjugate induced by pCS stimulation (10 µM, 24 h); data are mean ± s.e.m., n = 9 independent experiments. e) Pre-treatment of hCMEC/D3 cell monolayers with erlotinib (2.5 µM, 10 min) prevents the reduction in TEER induced by pCS stimulation (10 µM, 24 h); data are mean ± s.e.m., n = 9 independent experiments. f) Confocal microscopic analysis of expression of the tight junction component zona occludens-1 (ZO-1; white) in hCMEC/D3 cells following treatment with either pCS (10 µM, 24 h), erlotinib (2.5 µM, 24 h) or both (erlotinib 2.5 µM, 10 min pre-treatment, followed by pCS 10 µM, 24 h); nuclei are counterstained with DAPI (blue), scale bar = 15 µm. Images are representative of at least three independent experiments. g) Confocal microscopic analysis of expression of AF488-conjugated phalloidin labelled actin filaments (white) in hCMEC/D3 cells following treatment with either pCS (10 µM, 24 h), erlotinib (2.5 µM, 24 h) or both (erlotinib 2.5 µM, 10 min pre-treatment, followed by pCS 10 µM, 24 h); nuclei are counterstained with DAPI (blue), scale bar = 15 µm. Images are representative of at least three independent experiments. h) Paracellular permeability to a 70 kDa FITC-dextran conjugate of hCMEC/D3 cell monolayers bearing siRNA sequences targeting the EGFR (siRNA 5, siRNA 11, siRNA 12), a non-targeting negative control siRNA sequence, or that had been mock transfected, with or without stimulation with pCS (10 µM, 24 h); data are mean ± s.e.m., n = 4 independent experiments. i) TEER of hCMEC/D3 cell monolayers bearing siRNA sequences targeting the EGFR (siRNA 5, siRNA 11, siRNA 12), a non-targeting negative control siRNA sequence, or that had been mock transfected, with or without stimulation with pCS (10 µM, 24 h); data are mean ± s.e.m., n = 3 independent experiments.

Considering these interactions, we investigated whether EGFR activation mediated the permeabilising actions of pCS upon hCMEC/D3 cells. Pre-treatment with erlotinib (2.5 µM, 10 min pre-treatment) efficiently ablated the pCS-induced (10 µM, 24 h) increase in paracellular permeability (Fig. 3D) and decrease in TEER (Fig. 3E). These functional effects were paralleled by a reduction in pCS-stimulated disruption to the intracellular distribution of the tight junction molecule ZO-1 (Fig. 3F) and to the actin cytoskeleton (Fig. 3G).

While erlotinib has high selectivity for the EGFR, it is not without off-target effects; hence, we confirmed the centrality of this receptor in pCS action through use of targeted siRNA-mediated knockdown. Treatment of hCMEC/D3 cells expressing any of three independent siRNA sequences (knockdown efficiencies of 65.3 %, 56.7 % and 76.8 % respectively, Extended Data Fig. 5) with pCS (10 µM, 24 h) had little effect on either paracellular permeability (Fig. 3H) or TEER (Fig. 3I). This was in marked contrast to cells that had been either mock transfected or transfected with non-targeting siRNA negative control sequence, in which pCS exposure significantly (*P* < 0.05) increased paracellular permeability (Fig. 3H) and significantly (*P* < 0.05) decreased TEER (Fig. 3I), closely aligning with previous experiments.

### pCS stimulation of EGFR activates an ANXA1–STAT3–MMP signalling pathway

EGFR is coupled to a number of intracellular signalling pathways, but one potentially relevant target is the protein annexin A1 (ANXA1)^35^, a molecule we have previously identified as a major regulator of BBB function^36^. We therefore investigated whether this protein was involved in the response to pCS. Initial experiments confirmed that ANXA1 underwent Tyr21 phosphorylation in response to pCS exposure (10 µM, 30 min; Fig. 4A) in an erlotinib-sensitive manner (2.5 µM, 10 min pre-treatment; Fig. 4B). Using hCMEC/D3 cells stably bearing shRNA sequences to ANXA1^10^ (Extended Data Fig. 5), we further identified that exposure to pCS (10 µM, 24 h) significantly enhanced paracellular permeability in monolayers of wild-type and scramble shRNA-bearing hCMEC/D3 cells, but that this response was notably absent in cells transfected with ANXA1 shRNA sequences (Fig. 4C). Similarly, TEER was significantly (*P* < 0.01) reduced in wild-type and scramble shRNA-bearing hCMEC/D3 cells upon pCS exposure, but not in ANXA1 shRNA transfected cells (Fig. 4D).

**Figure 4:**
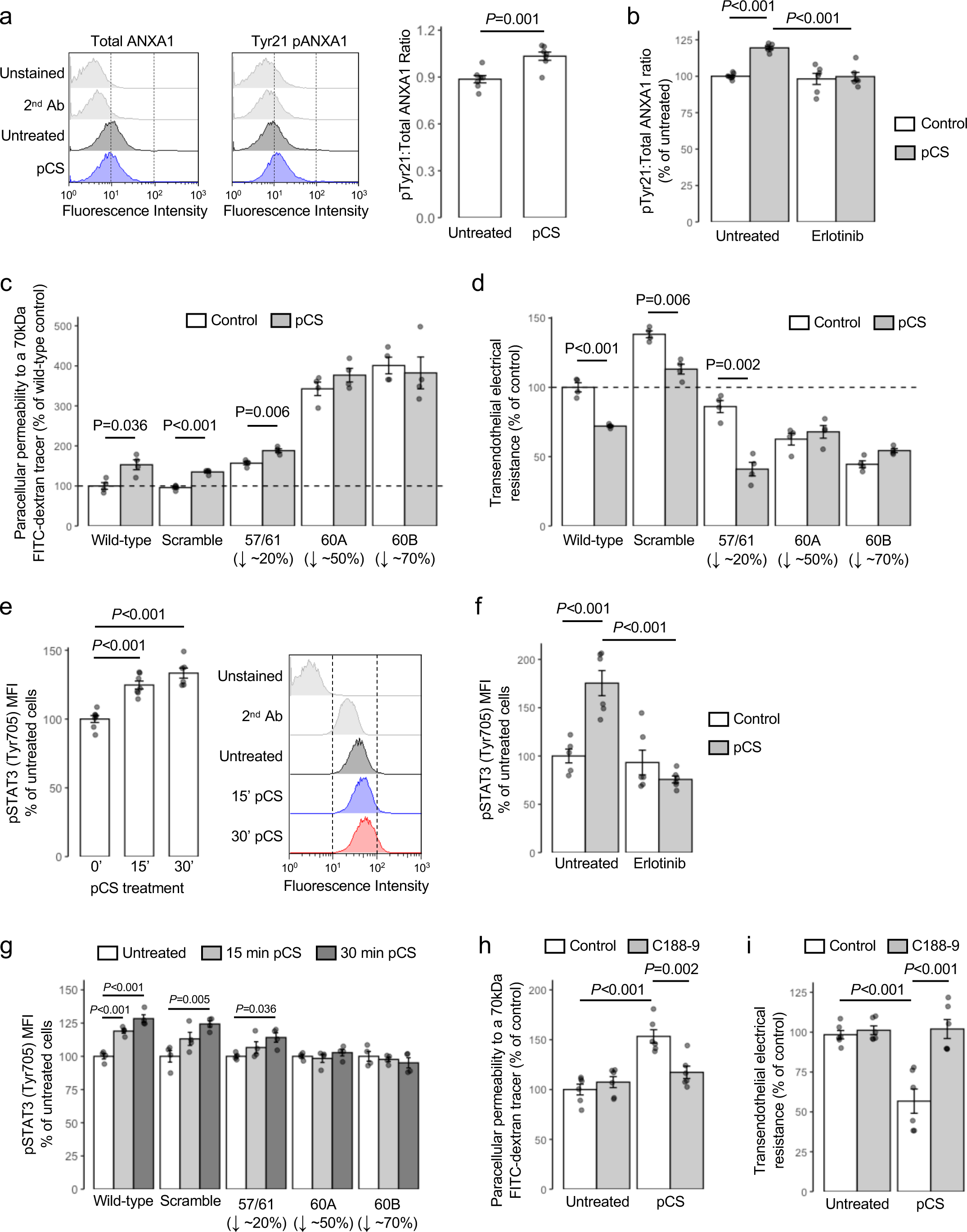
Stimulation of EGFR by pCS activates a downstream ANXA1–STAT3 signalling pathway. a) Stimulation of hCMEC/D3 cells with pCS (10 µM, 30 min) increases the ratio of Tyr21 phosphorylated to total annexin A1; data are mean ± s.e.m., n = 7 independent experiments; representative flow cytometry histograms are shown. b) Pre- treatment of hCMEC/D3 cells with erlotinib (2.5 µM, 10 min) prevents the increase in the Tyr21 phosphorylated to total ANXA1 expression ratio induced by pCS stimulation (10 µM, 30 min); data are mean ± s.e.m., n = 6 independent experiments. c) Paracellular permeability to a 70 kDa FITC-dextran tracer of monolayers of wild-type hCMEC/D3 cells, or hCMEC/D3 cells stably transfected with either a scramble shRNA sequence or one of three independent shRNA sequences targeting ANXA1 following stimulation with pCS (10 µM, 24 h); data are mean ± s.e.m., *n* = 4 independent experiments. d) TEER of monolayers of wild-type hCMEC/D3 cells, or hCMEC/D3 cells stably transfected with either a scramble shRNA sequence or one of three independent shRNA sequences targeting ANXA1 following stimulation with pCS (10 µM, 24 h); data are mean ± s.e.m., *n* = 4 independent experiments. e) Stimulation of hCMEC/D3 cells with pCS (10 µM) causes a time-dependent increase in expression of Tyr705 phosphorylated STAT3; data are mean ± s.e.m., n = 7 independent experiments; representative flow cytometry histograms are shown. f) Pre-treatment of hCMEC/D3 cells with erlotinib (2.5 µM, 10 min) prevents the increase in Tyr705 phosphorylated STAT3 expression induced by pCS treatment (10 µM, 30 min); data are mean ± s.e.m., n = 6 independent experiments. g) Expression of Tyr705 phosphorylated STAT3 in wild-type hCMEC/D3 cells, or hCMEC/D3 cells stably transfected with either a scramble shRNA sequence or one of three independent shRNA sequences targeting ANXA1 following stimulation with pCS (10 µM), expressed as percentage of untreated cells; data are mean ± s.e.m., n = 4 independent experiments. h) Pre-treatment of hCMEC/D3 cell monolayers with the STAT3 inhibitor C188-9 (5 µM, 30 min) prevents the increase in paracellular permeability to a 70 kDa FITC-dextran conjugate induced by pCS stimulation (10 µM, 24 h); data are mean ± s.e.m., n = 6 independent experiments. i) Pre- treatment of hCMEC/D3 cell monolayers with C188-9 (5 µM, 30 min) prevents the reduction in TEER induced by pCS stimulation (10 µM, 24 h); data are mean ± s.e.m., n = 6 independent experiments.

Both EGFR signalling and ANXA1 Tyr21 phosphorylation have been linked with the transcription factor STAT3^37,38^, hence we examined the impact of pCS exposure upon STAT3 activation. Exposure of hCMEC/D3 cells to pCS (10 µM) significantly (*P* < 0.001) enhanced STAT3 Tyr705 phosphorylation within 15 min (Fig. 4E), an effect that was not seen in cells pre-treated with erlotinib (2.5 µM, 10 min pre-treatment; Fig. 4F) and was attenuated or absent in hCMEC/D3 cells bearing ANXA1 shRNA sequences (Fig. 4G). Notably, pre-treatment of hCMEC/D3 cells with the selective STAT3 inhibitor C188-9 (5 µM, 30 min pre-treatment) significantly (*P* ≤ 0.002) reduced the effects of pCS exposure (10 µM, 24 h) upon both paracellular permeability to a 70 kDa FITC-dextran tracer (Fig. 4H) and TEER (Fig. 4I).

### Inhibition of MMP2/9 activity prevents the permeabilising effects of pCS in vitro

Activation of MMPs, particularly that of MMP-2 and MMP-9, is known to be downstream of EGFR^34^ and STAT3^39^ activation, to be capable of damaging BBB permeability^40^, and to be associated with CKD^41^, leading us to hypothesise that the permeabilising effects of pCS might be mediated through these enzymes. Stimulation of hCMEC/D3 cells with pCS (10 µM, 24 h) significantly (*P* ≤ 0.001) increased MMP-2 and MMP-9 expression (Fig. 5A), a response inhibited by pre-treatment with either the EGFR inhibitor erlotinib (2.5 µM, 10 min pre-treatment) or the STAT3 inhibitor C188-9 (5 µM, 30 min pre-treatment; Fig. 5B). Analysis of hCMEC/D3 conditioned medium by gelatin zymography confirmed that these pCS-induced (10 µM, 24 h) increases in cellular MMP-2 and MMP-9 expression were mirrored by enhanced gelatinase activity (Fig. 5C), an increase again sensitive to erlotinib pre-treatment (2.5 µM, 30 min prior to 10 µM pCS; Fig. 6D).

**Figure 5:**
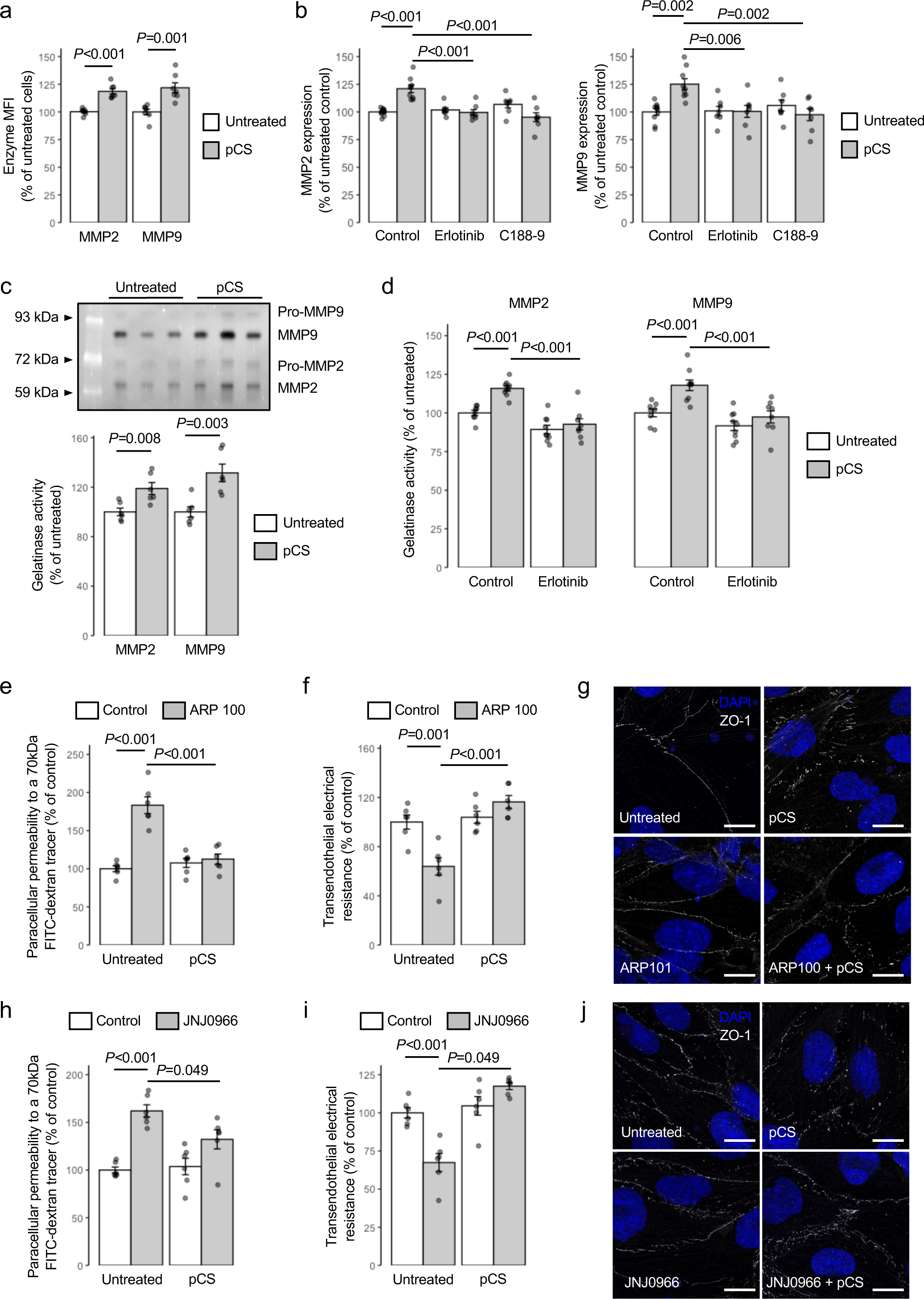
pCS enhances endothelial permeability *via* mobilisation of active MMP2 and MMP9. a) Treatment of hCMEC/D3 cells with pCS (10 µM, 24 h) increases expression of MMP2 and MMP9; data are mean ± s.e.m., n = 7 independent experiments. b) Stimulation of MMP2 or MMP9 expression in hCMEC/D3 cells by pCS (10 µM, 24 h) is prevented by either erlotinib (2.5 µM, 10 min pre-treatment) or C188-9 (5 µM, 30 min pre-treatment); data are mean ± s.e.m., n = 6 independent experiments. c) Active MMP2 and MMP9 are secreted into the culture medium of hCMEC/D3 cells in response to pCS (10 µM, 24 h) stimulation; data are mean ± s.e.m., n = 6 independent experiments; a typical gelatin zymography is shown, highlighting both active MMP2 and MMP9 and artifactually revealed pro-MMP2 and pro-MMP9 activity. d) Active MMP2 and MMP9 release from hCMEC/D3 cells into culture medium in response to pCS (10 µM, 24 h) stimulation is prevented by erlotinib (2.5 µM, 10 min pre- treatment); data are mean ± s.e.m., n = 11-12 independent replicates. e) Pre-treatment of hCMEC/D3 cell monolayers with the MMP2 inhibitor ARP 100 (12 nM, 15 min) prevents the increase in paracellular permeability to a 70 kDa FITC-dextran conjugate induced by pCS stimulation (10 µM, 24 h); data are mean ± s.e.m., n = 6 independent experiments. f) Pre- treatment of hCMEC/D3 cell monolayers with ARP 100 (12 nM, 15 min) prevents the reduction in TEER induced by pCS stimulation (10 µM, 24 h); data are mean ± s.e.m., n = 6 independent experiments. g) Confocal microscopic analysis of expression of the tight junction component zona occludens-1 (ZO-1; white) in hCMEC/D3 cells following treatment with either pCS (10 µM, 24 h), ARP 100 (12 nM, 24 h) or both (ARP 100 12 nM, 15 min pre-treatment, followed by pCS 10 µM, 24 h); nuclei are counterstained with DAPI (blue), scale = 15 µm . Images are representative of at least three independent experiments. h) Pre-treatment of hCMEC/D3 cell monolayers with the MMP9 inhibitor JNJ0966 (440 nM, 15 min) prevents the increase in paracellular permeability to a 70 kDa FITC-dextran conjugate induced by pCS stimulation (10 µM, 24 h); data are mean ± s.e.m., n = 6 independent experiments. i) Pre-treatment of hCMEC/D3 cell monolayers with JNJ0966 (440 nM, 15 min) prevents the reduction in TEER induced by pCS stimulation (10 µM, 24 h); data are mean ± s.e.m., n = 6 independent experiments. j) Confocal microscopic analysis of expression of the tight junction component zona occludens-1 (ZO-1; white) in hCMEC/D3 cells following treatment with either pCS (10 µM, 24 h), JNJ0966 (440 nM, 24 h) or both (JNJ0966 440 nM, 15 min pre-treatment, followed by pCS 10 µM, 24 h); nuclei are counterstained with DAPI (blue), scale = 15 µm. Images are representative of at least three independent experiments.

**Figure 6:**
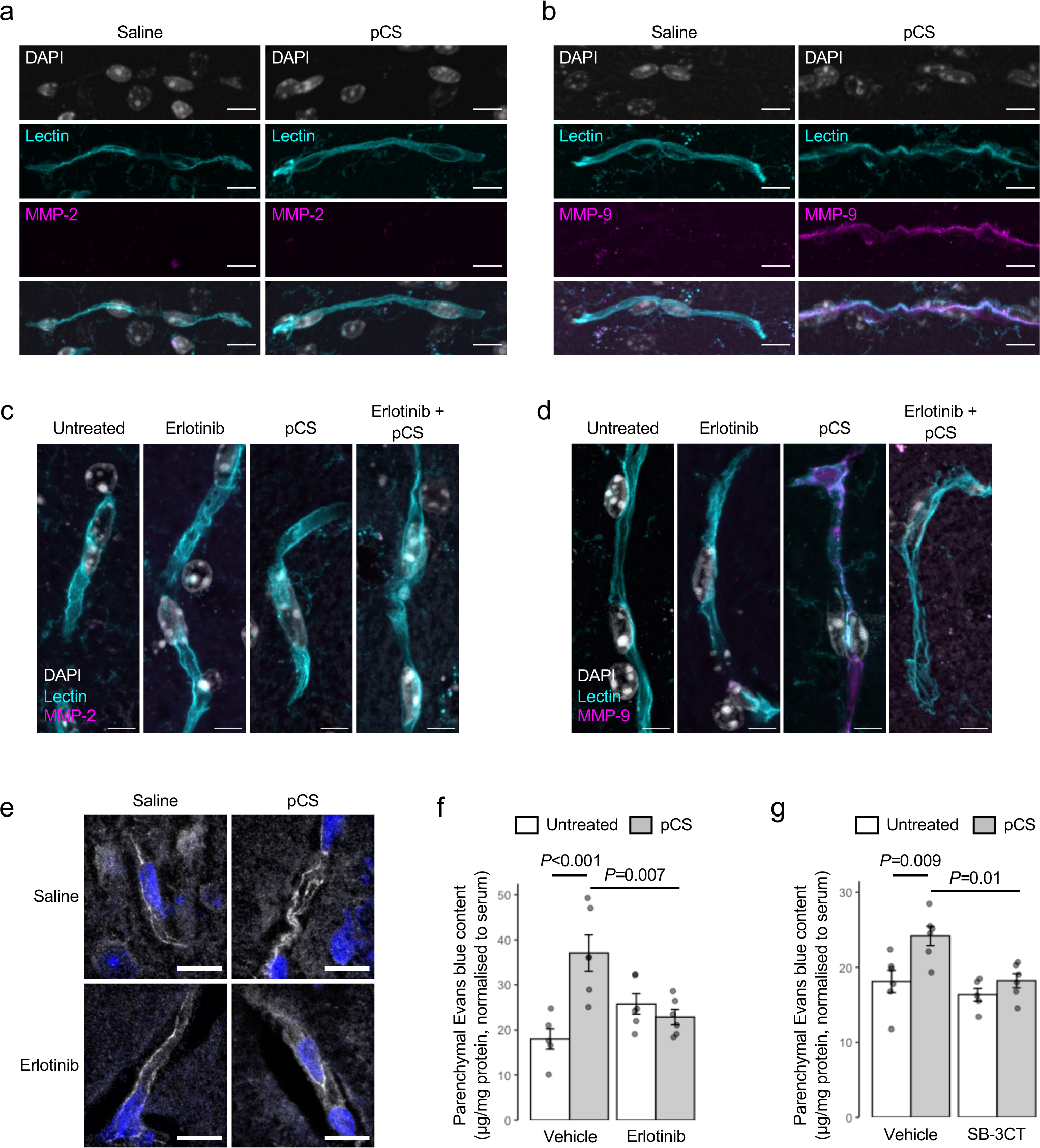
Treatment with pCS induces BBB permeability *in vivo* through EGFR- dependent mobilisation of MMP9. a, b) Confocal microscopic analysis of expression of a) MMP-2 or b) MMP-9 (magenta) in the brains of male C57Bl/6 exposed to pCS by s.c. implantation with osmotic mini-pumps releasing at 7.5 µg/h for 28 days, blood vessels are defined by tomato lectin binding (cyan), nuclei are counterstained with DAPI (white), scale = 10 µm, images are representative of 7-10 animals. c, d) Confocal microscopic analysis of c) MMP2 and d) MMP9 expression (magenta) within the cerebral microvasculature of male C57Bl/6 mice exposed to pCS (10 mg/kg, i.p. 2 h) with or without erlotinib pre-treatment (50 mg/kg, i.p. 1 h pre-treatment); blood vessels are defined by tomato lectin binding (cyan), nuclei are counterstained with DAPI (white), scale bar = 5 µm, images are representative of 7-10 animals. e) Typical expression of ZO-1 (white) within the cerebral microvasculature of male C57Bl/6 mice exposed to pCS (10 mg/kg, i.p. 2 h) with or without erlotinib pre-treatment (50 mg/kg, i.p. 1 h pre-treatment); nuclei are counterstained with DAPI (blue), scale bar = 10 µm. Typical expression of ZO-1 (white) within the cerebral microvasculature of male C57Bl/6 mice exposed to pCS (10 mg/kg, i.p. 2 h) with or without erlotinib pre-treatment (50 mg/kg, i.p. 1 h pre-treatment); nuclei are counterstained with DAPI (blue), scale bar = 10 µm. f) Increased extravasation of Evans blue tracer into the parenchyma of male C57Bl/6 mice following i.p. injection of pCS (10 mg/kg, 2 h exposure) is prevented by erlotinib administration (50 mg/kg i.p., 1 h pre-treatment); data are mean ± s.e.m., n = 5-6 animals. g) Increased extravasation of Evans blue tracer into the parenchyma of male C57Bl/6 mice following i.p. injection of pCS (10 mg/kg, 2 h exposure) is prevented by administration of the dual-specific MMP2/MMP9 inhibitor SB-3CT (10 mg/kg i.p., 1 h pre-treatment); data are mean ± s.e.m., n = 5-6 animals.

To judge the relevance of this increase in MMP activity to endothelial barrier integrity, we investigated the effect of selective MMP inhibitors upon the actions of pCS. Pre-treatment of hCMEC/D3 monolayers with either the selective MMP-2 inhibitor ARP 100 (12 nM, 15 min pre-treatment) or the selective MMP-9 inhibitor JNJ0966 (440 nM, 15 min pre-treatment) prevented the disruptive effects of pCS exposure (10 µM, 24 h) upon paracellular permeability to a 70kDa FITC-dextran tracer (Fig. 5E, H), TEER (Fig. 5F, I) and intracellular distribution of the tight junction anchor ZO-1 (Fig. 5G, J).

### pCS impairs BBB integrity through an EGFR–STAT3–MMP-9 pathway in vivo

Having established the EGFR–ANXA1–STAT3–MMP-2/9 pathway as mediating the effects of pCS upon endothelial cell barrier function *in vitro*, we sought to validate this pathway *in vivo*. Examination of C57Bl/6 male mice exposed to elevated circulating pCS for 4 weeks (Extended Data Table 3) revealed a clear increase in brain blood vessel-associated MMP-9 (though not MMP-2) expression (Fig. 6A-B), confirming that at least a similar pathway to that seen in human cells is operative in mice. Furthermore, pre-treatment of male C57Bl/6 mice with the EGFR inhibitor erlotinib (50 mg/kg, i.p., 1 h pre-treatment) prior to pCS injection (10 mg/kg i.p., 2 h) suppressed the pCS-induced immunohistochemical appearance of Tyr21 phospho- ANXA1 (Extended Data Fig. 6) and MMP-9 (Fig. 6C-D) within the cerebrovascular wall, ameliorated pCS-induced disruption of vascular ZO-1 expression (Fig. 6E) and prevented a pCS-induced increase in parenchymal Evans blue tracer extravasation (Fig. 6F). Confirming the importance of pCS-induced MMP activity in mediating BBB disruption, pre-treatment of mice with the MMP-2/9 dual-specific inhibitor SB-3CT (10 mg/kg i.p., 1 h pre-treatment) also prevented pCS treatment (10 mg/kg i.p., 2h) from increasing parenchymal Evans blue extravasation (Fig. 6G).

### Treatment with the EGFR inhibitor erlotinib can prevent the permeabilising effect of CKD patient serum upon hCMEC/D3 monolayers

While kidney failure is associated with damage to the cerebral vasculature, it is also associated with altered circulating levels of numerous uraemic retention molecules, of which pCS is only one, albeit important, example. Most notably, the EGFR has been demonstrated to interact with the gut microbe-derived uraemic toxin indoxyl sulfate (IS)^42^, although treatment of hCMEC/D3 monolayers with this metabolite was without effect on paracellular permeability to a 70 kDa FITC-dextran tracer or TEER at any concentration tested (5 µM, 25 µM, 50 µM, 150 µM; Extended Data Fig. 7).

To investigate the relative importance of pCS in mediating the cerebrovascular symptoms of CKD, we exposed hCMEC/D3 monolayers for 24 h to medium containing 20 % de- complemented serum from individuals undergoing haemodialysis or age-matched individuals with normal kidney function who were undergoing evaluation as potential kidney donors (full patient details in Table 2), either alone or in combination with erlotinib (2.5 µM, 10 min pre-treatment). Treatment of hCMEC/D3 cells with haemodialysis patient sera, but not with that from healthy donors, significantly (*P* = 0.004) enhanced paracellular permeability to a 70 kDa FITC-dextran tracer, an effect reversed by erlotinib pre-treatment (Fig. 7A), although notably none of these treatments affected TEER (Fig. 7B). Additionally, similar treatment of hCMEC/D3 cells with sera from haemodialysis patients, but again not with that from healthy donors, significantly (*P* < 0.05) increased intracellular expression of both MMP-2 (Fig. 7C) and MMP-9 (Fig. 7D).

**Figure 7:**
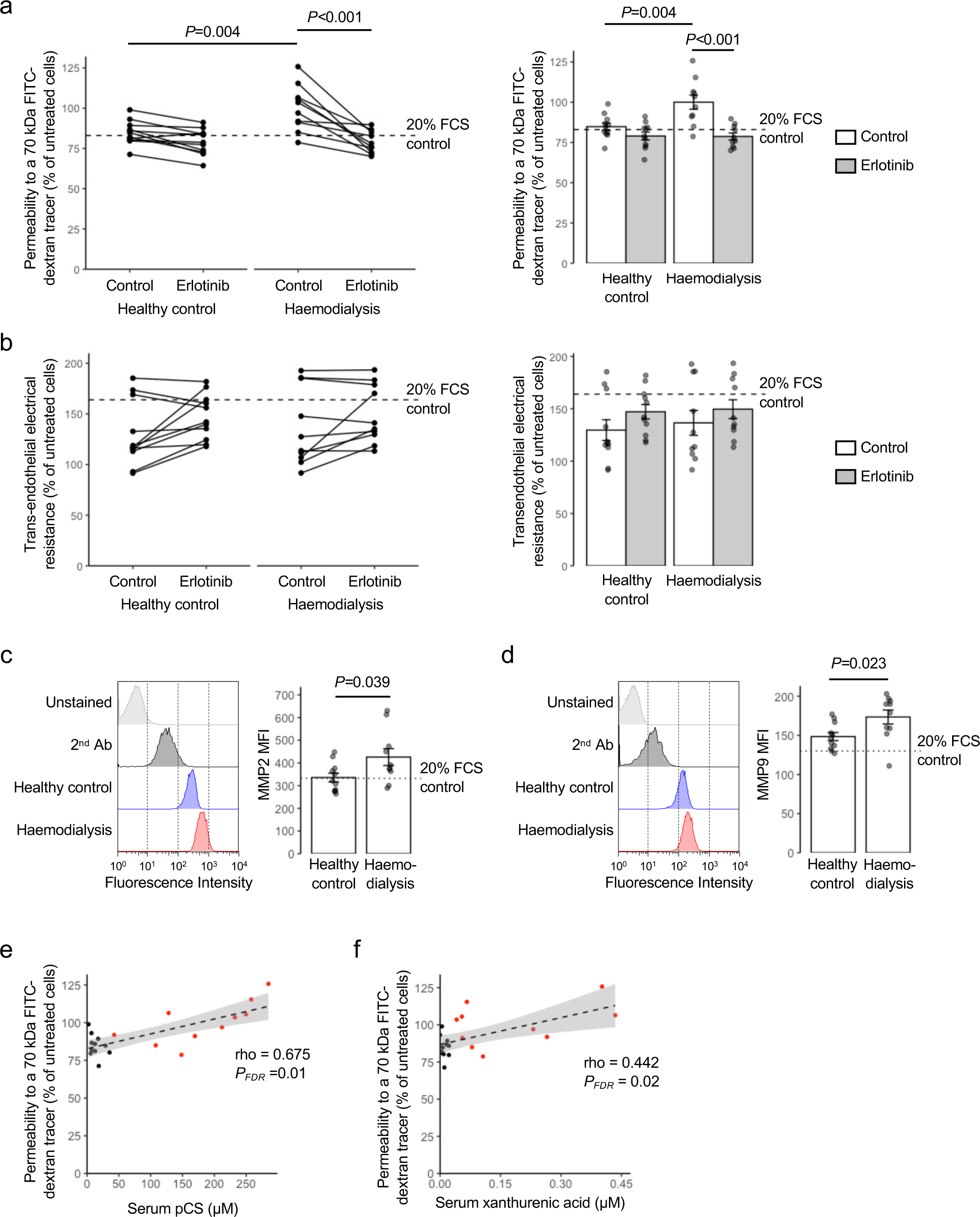
Sera from human haemodialysis patients significantly enhances endothelial permeability, an effect prevented by erlotinib pre-treatment. a) Treatment of hCMEC/D3 monolayers with medium containing de-complemented sera from patients undergoing haemodialysis (20 %, 24 h) significantly enhances permeability to a 70 kDa FITC-dextran tracer, an effect not seen with equivalent sera from healthy donors, and prevented by inclusion of erlotinib (2.5 µM, 10 min); data are mean ± s.e.m., n = 10-11 donors. b) TEER of hCMEC/D3 monolayers is not affected by treatment with media containing sera (20 %, 24 h) from either haemodialysis patients or healthy donors, nor by erlotinib inclusion (2.5 µM, 10 min pre- treatment); data are mean ± s.e.m., n = 10-11 donors. c) Treatment of hCMEC/D3 cells with medium containing de-complemented sera (20 %, 24 h) from patients undergoing haemodialysis, but not with that from healthy donors, increases expression of MMP2; data are mean ± s.e.m., n = 10-11 donors. d) Treatment of hCMEC/D3 cells with medium containing de-complemented sera (20 %, 24 h) from patients undergoing haemodialysis, but not with that from healthy donors, increases expression of MMP9; data are mean ± s.e.m., n = 10-11 donors. e) Correlation between concentrations of pCS in human sera samples (healthy donors in black, haemodialysis patients in red) with 70 kDa FITC-dextran permeability data from (a) above; n = 10-11 donors. f) Correlation between concentrations of xanthurenic acid in human sera samples (healthy donors in black, haemodialysis patients in red) with 70 kDa FITC- dextran permeability data from (a) above; n = 10-11 donors.

Having established that sera from individuals with kidney failure could modulate permeability of our *in vitro* BBB model in an erlotinib-sensitive manner, we examined the relative contributions of different amino acid-derived potential uraemic toxins to this response. Following analysis of serum concentrations of different metabolites by LC-MS/MS (Table 3; mean pCS concentrations: healthy donor 14.12 ± 3.09 µM, haemodialysis patient 155.26 ± 27.36 µM, *P*<0.001), we correlated these with changes in paracellular permeability; only correlations with pCS (rho = 0.675, *P*FDR = 0.01; Fig. 7E) and the tryptophan metabolite xanthurenic acid (rho = 0.442, *P*FDR = 0.02; Fig. 7F) remained significant following correction for multiple testing.

## Discussion

Impaired kidney function has serious consequences for virtually every organ system in the body, and the central nervous system is no exception. Patients with CKD, including those at not yet requiring dialysis, exhibit significant cognitive decline^43^ and enhanced stroke risk^44^, even after correction for typical cardiovascular risk factors. Cerebral microvascular disease and BBB disruption are thought to underlie these links^45^, but little is known about how renal impairment results in BBB damage. Here we present data indicating a novel BBB-disruptive role for the major uraemic toxin and gut microbe-host co-metabolite pCS, acting *via* the EGFR to mobilise MMP activity. Moreover, we show that pCS treatment induces widespread transcriptomic changes within the brain, rapidly dampening numerous aspects of neuronal activity and metabolism. Together, these data strongly implicate pCS as a major contributory factor to the links between kidney failure and impaired brain function.

Interactions of pCS with the peripheral vasculature have previously been reported, with evidence of arterial leakage in patients with CKD^46^ and in animal^28^ and *in vitro* models of uraemia^47^. Mechanistically, pCS at concentrations found in uraemia can cause inter-endothelial adherens junction disruption^46^, potentially underlying uraemia-associated endothelial leakage. Here, we extend the damaging actions of pCS to the cerebral vasculature and BBB *in vivo*, acting at concentrations similar to those of free pCS seen at stages III-IV of CKD^48^. We further provide mechanistic evidence coupling pCS agonism at the EGFR, *via* phosphorylation of ANXA1 and STAT3 to mobilisation of MMP activity, a known contributor to neurovascular disruption^49^. Importantly, these actions appear clinically relevant, with the serum of human haemodialysis patients impairing *in vitro* brain endothelial permeability barrier with a potency highly correlated to levels of pCS, an effect prevented by co-administration of the EGFR antagonist erlotinib. This positioning of the EGFR as a major mediator of pCS actions in the brain may additionally point to potential therapeutic strategies guarding against cerebrovascular damage in CKD; several EGFR inhibitors (e.g., erlotinib, gefitinib) and monoclonal antibodies (e.g., cetuximab, necitumumab) are licenced for clinical use in malignant disease, suggesting there may be scope for broader use in CKD, although caution is needed regarding the safety of at least some of these agents in advanced renal impairment^50^. Nonetheless, this highlights the possibility of developing an alternative risk-reduction strategy for cerebrovascular complications in CKD, something currently chiefly addressed through management of hypertension.

Cerebral small vessel disease, for which BBB failure is a fundamental driver^51^, is thought to contribute to both the cognitive impairments and enhanced stroke risk seen in CKD patients^52^. What is less clear, partly due to the obscuring effects of the relationship between CKD and cerebrovascular risk factors such as diabetes and hypertension, is why this is the case. Activation of MMPs within the neurovascular unit has been reported in several forms of cerebrovascular disease, including stroke^53^, traumatic injury^54^, cerebral small vessel disease^55^ and, notably, in murine models of CKD^56^. By elaborating a mechanistic link between gut microbe/host co-derived pCS, cerebrovascular MMP activity and impaired BBB integrity, we provide an explanation for how cerebral vessels may be damaged in CKD. Moreover, whilst local MMP activity has been demonstrated in a range of kidney disorders^57^, whether this holds true in the brains of CKD patients is unknown. Our data indicating both a close correlation between pCS content and the BBB-permeabilising actions of haemodialysis patient serum, and moreover that exposure to such serum directly enhances endothelial MMP expression, suggest that this is highly likely.

While enhanced BBB permeability is a recognised driver of damage to cognitive function^58^, our data suggest that the effects of pCS upon brain function extend beyond its immediate impact on the cerebral vasculature. Transcriptional analysis of pCS-exposed mice indicates a more generalised inhibitory effect of the toxin upon neural activity, which might also be expected to impair cognitive abilities. We did not detect such cognitive disruption following long-term pCS exposure however, a finding with several potential explanations. Previous studies identifying cognitive sequelae of BBB damage have shown a substantial increase in permeability, in the order of a 5 fold increase or greater^59,60^, to be associated with cognitive deficits. In our study, animals exposed for one month to pCS exhibited an approximately 30% increase in tracer permeability across the BBB; it may be that the degree of permeability deficit caused in this study was insufficient to impair cognition to a degree detectable using the tests employed. Notably, previous studies employing higher pCS concentrations have shown the metabolite to impair cognition in both humans^61^ and mice^59^.

Intriguingly, while pCS clearly influenced BBB permeability, we found no evidence that this was true for the other major gut microbe-derived uraemic toxin, IS. This is perhaps surprising, given previous evidence that IS concentrations correlate with BBB impairment in partially nephrectomised ratsz^59^ and vascular disease in CKD patients^62^. In contrast, however, suppression of raised circulating IS by sequestration did not reverse adenine-rich diet induced BBB impairment^59^, highlighting the complexity of the pathways impinging on the BBB. Taken alongside our recent demonstration that the CKD-associated molecule trimethylamine *N*-oxide (TMAO) exerted marked beneficial effects upon both the BBB and cognition at levels seen in metabolically healthy individuals^10^, it appears the generalised assumption that uraemia- associated molecules are always deleterious may need revisiting^63^.

While pCS predominates in the circulation, it is not the only *p*-cresol conjugate found in blood, with pCG representing a significant, albeit minor, fraction^15^. Moreover, levels of pCG do increase markedly with worsening CKD stages, and have been shown to correlate with mortality risk in a similar way to pCS^64^. However, we have shown pCG to exert very different actions upon the BBB to pCS, with its administration at physiologically relevant levels *in vivo* enhancing barrier integrity by antagonism of basal circulating lipopolysaccharide interactions with TLR4^65^. Moreover, whole brain RNAseq analysis of pCG-exposed mice indicated a positive regulation in neuronal activity gene ontologies, notably opposite to the suppressive effects of pCS described here^65^. That the two major conjugates of gut microbe-derived *p*-cresol should so markedly differ in effect upon the brain is striking, emphasising the need for mechanistic studies of gut microbe/host co-metabolite action before conclusions can safely be drawn about their clinical role(s).

Most circulating pCS is ultimately derived from the actions of gut microbes upon dietary amino acids^11^, thus our description of its effects upon the BBB further emphasises the importance of this facet of gut microbe-host interactions. Notably, while we and others have shown other gut microbe-derived metabolites, including short-chain fatty acids^8,9^, methylamines^10^, and pCG^65^ to reinforce BBB integrity in healthy animals, pCS is overtly detrimental in its effects. Given the wide range of metabolites produced by gut microbes, that some agents are deleterious at supraphysiological concentrations is unsurprising, but does emphasise the importance of mechanistic analyses of gut-brain communication pathways. Overall, our description of pCS as a major link between gut microbes, kidney function and the cerebral vasculature highlights the role of the gut microbiota in at least some of the systemic effects of renal disease, emphasising its potential as a therapeutic target for clinical management of CKD.

## Methods

### Human serum samples

Pre-dialysis serum samples were collected from patients undergoing haemodialysis at Barts Health NHS Trust and from age-/sex-matched healthy controls undergoing evaluation for kidney donation who had previously consented for sample inclusion in the Barts Diabetic Kidney Disease Biobank, see Table 2 for demographic details. Ethical approval for the inclusion in the Barts DKC Biobank was provided under REC reference no. 18/EE/0142. Approval for the current study was provided under REC reference no. 20/LO/0361. The study followed established standard operating procedures and all experiments involving human tissue conformed to the principles set out in the World Medical Association Declaration of Helsinki. All serum samples were de-complemented by incubation at 56 °C for 20 min prior to use.

### Animals

Wild-type, male, specific pathogen-free, C57Bl/6J mice (Charles River UK Ltd., Margate, UK) were used for all experiments, aged between 7 and 8 weeks at the start of experiments. All experiments were approved by the QMUL Animal Welfare and Ethical Review Board (acute studies) or the University of East Anglia Animal Welfare and Ethical Review Body (long-term studies) and were performed in accordance with the ARRIVE guidelines and the UK Animals (Scientific Procedures) Act, 1986, under UK Home Office Project Licences PFA5C4F4F and PP9417531. Individual animals were used as experimental units; animals were randomly assigned to groups using the =rand() function of Microsoft Excel.

### Animals: acute studies

Mice were housed in groups of five per cage in a room with controlled lighting (lights on 08.00– 20.00 h) and temperature (21–23 °C) and with standard lab chow (5058, LabDiet, St. Louis, USA) and water available *ad libitum.* Mice were acclimatised to the animal facility environment for one week prior to experimentation, randomised to treatment group, treated as described below, and killed by transcardial perfusion with ice-cold 0.1 M phosphate buffered saline (PBS) under deep sodium pentobarbital anaesthesia (50 mg/kg, i.p.). Whole blood was collected and permitted to coagulate for 30 minutes at room temperature, prior to centrifugation at 800g for separation of serum, and brains were rapidly removed and bisected. For immunohistochemical analysis, half-brains were fixed by immersion in 4 % formaldehyde in 0.1 M PBS at 4 °C for 24 h, after which they were embedded in paraffin wax. For RNAseq studies, half-brains were stored in RNAlater (ThermoFisher Scientific Ltd., UK) at 4 °C overnight, and then frozen at -80 °C for later RNA extraction as described below. For studies of Evans blue dye extravasation, half-brains were collected into 0.1 M phosphate buffered saline (PBS) at 4 °C for later analysis as described below. In all cases, further tissue analysis was performed blinded to original groupings.

### Animals: long-term studies

Mice were housed in groups of four per cage in a room with controlled lighting (lights on 08.00– 20.00 h) and temperature (21–23 °C) and with standard lab chow (5058, LabDiet, St. Louis, USA) and water available *ad libitum*. Mice were acclimatised to the animal facility environment for two weeks prior to experimentation. Animals were anaesthetised by isofluorane inhalation and subcutaneously implanted with osmotic minipumps (model 2004, Alzet Cupertino, CA, USA) previously loaded with either 30 mg/ml pCS or saline vehicle, designed to release contents at 0.25 µl/h for a total period of 4 weeks, with minipump contents assigned randomly across groups. At the end of this period mice were killed by transcardial perfusion with ice-cold 0.1 M phosphate buffered saline (PBS) under deep sodium pentobarbital anaesthesia (50 mg/kg, i.p.). Whole blood was collected and permitted to coagulate for 30 minutes at room temperature, prior to centrifugation at 800g for separation of serum, and brains were rapidly removed and bisected. For immunohistochemical analysis, half-brains were fixed by immersion in 4 % formaldehyde in 0.1 M PBS at 4 °C for 24 h, after which they were embedded in paraffin wax. For studies of sodium fluorescein extravasation, half-brains were collected into 0.1 M phosphate buffered saline (PBS) at 4 °C for later analysis as described above. In all cases, further tissue analysis was performed blinded to original groupings.

### In vivo BBB permeability analysis

Mice were injected i.p. with 100 µl of a 2 % w/v solution of Evans blue tracer in 0.9 % saline (Merck Ltd, Poole, UK) or i.v. with 200 µl of a 2% w/v solution of sodium fluorescein in 0.9 % saline (Merck Ltd., UK). Tracer was permitted to circulate for 1 h before animals were deeply anaesthetised with sodium pentobarbital (50 mg/kg, i.p.) and, following sampling of blood by cardiac puncture, transcardially perfused with 0.9 % PBS at 4 °C to remove dye remaining within the vasculature. Brains were removed, bisected and left hemispheres were homogenised by maceration in 0.1 M PBS. Suspended macromolecules were precipitated by incubation with 60 % trichloroacetic acid, and tracer content of the resulting supernatants was assessed using a CLARIOstar fluorescence spectrophotometer, set to read absorbance at 620 nm for Evans blue, or fluorescence at 516 nm for sodium fluorescein (BMG Labtech GmbH, Germany) alongside a standard curve of defined concentrations of tracer in the same buffer. Brain tracer content was expressed as µg of dye/mg of brain tissue, normalised to circulating serum concentrations.

### RNAseq analysis

Mice were transcardially perfused with 0.9 % saline at 4 °C to remove circulating blood, and brains were removed and collected into RNAlater (Thermofisher Scientific Ltd., UK) prior to storage at -20 °C for later analysis. Whole brain total RNA was extracted using a PureLink RNA Mini Kit (Thermofisher Scientific Ltd., UK) and quantified using a CLARIOstar spectrophotometer equipped with an LVis microplate (BMG Labtech GmbH, Germany). RNA samples (*n* = 5 pCS, *n* = 5 saline control) were sent to Macrogen Inc. (Republic of Korea) where they were subject to quality checks (RIN analysis); libraries were prepared (TruSeq Stranded mRNA LT Sample Prep Kit) for paired-end (2× 100 nt) sequencing on an Illumina HiSeq 4000 apparatus. Raw RNAseq sequence data (delivered in fastq format) were processed in-house as follows. Single-end forward reads were mapped onto the mouse genome (mm10) using HISAT2 v2.2.1^66^. The number of reads in each sample that mapped to exons in the BAM files returned by HISAT2 was determined using featureCounts within subread v2.0.1^67^. DESeq2 v1.34.0^68^ was used to identify significantly (P < 0.05, Benjamini-Hochberg) differentially expressed genes in the dataset. Entrez gene identifiers were converted to gene symbols using *Mus musculus* annotations downloaded from NCBI on 12 October 2022. Enrichr^33^ was used to perform Gene Ontology analysis. SPIA v2.48.0^69^ was used to identify KEGG pathways inhibited or activated in mouse brain samples exposed to pCS. KEGGgraph v1.56.0^70^ was used to generate the topological network from the log_2_ fold change values for the significantly differentially expressed genes and KEGG xml files for the *Mus musculus* pathways of interest. igraph v1.3.5^71^ was used to generate summary network statistics. Raw RNAseq data have been deposited with ArrayExpress under accession number E-MTAB-12326.

### Immunohistochemistry

Paraffin-embedded brains were sectioned (5 µm) using a rotary microtome and collected onto glass microscope slides. Following deparaffinisation with xylene and rehydration using graded ethanol:water solutions, heat-mediated antigen retrieval was performed by incubation in 10 mM Tris base, 1 mM EDTA, 0.05 % Tween-20, pH 9.0 at 90 °C for 20 min. Once cooled, endogenous peroxide activity was quenched by incubation for 15 min in 0.3 % H_2_O_2_ in Tris-buffered saline (TBS; 50 mM Tris base, 150 mM NaCl, pH 7.4). Sections were permeabilised by incubation in TBS containing 0.025 % Triton X-100 for 10 min, blocked by incubation in 10 % normal goat serum in TBS for 30 min and treated with primary antibodies overnight at 4 °C (MMP2: rabbit anti-human, 1:1000 Insight Biotechnology Ltd., UK, MMP9: mouse anti-human, 1:50, Santa Cruz Biotechnology Inc, USA, phosphotyrosine-21 ANXA1: rabbit anti-human, 1:1000^72^, ZO-1: rabbit anti-human 1:100, Thermofisher Scientific, UK), diluted in TBS containing 1 % normal goat serum, 0.025 % triton X-100, pH 7.4). Sections were washed thoroughly with TBS containing 1 % normal goat serum and incubated with relevant secondary antibody for 1 h at room temperature (horseradish peroxidase-conjugated goat anti-rabbit or anti-mouse 1:500, both Stratech Scientific, UK, AF488-conjugated goat anti-rabbit or anti-mouse 1:500, both Thermofisher Scientific, UK; diluted in TBS containing 1 % normal goat serum, 0.025 % triton X-100, pH 7.4). Sections were thoroughly washed in TBS and where appropriate peroxidase staining was developed using 0.5 mg/ml diaminobenzidine hydrochloride and 0.01 % H_2_O_2_. Sections were counterstained with haematoxylin or 50 ng/ml 4′,6-diamidino-2-phenylindole (DAPI) as appropriate and mounted for microscopic examination. Brightfield images were captured using a using a Nikon Eclipse 80i Stereology Microscope fitted with an Optronics Camera, using a 40x objective (NA: 0.75 mm, working distance: 0.66 mm). Fluorescence images were captured with an LSM880 confocal laser scanning microscope (Carl Zeiss Ltd., Cambridge, UK) fitted with 405, 488 and 561 nm lasers and an ×63 oil immersion objective lens (NA: 1.4 mm, working distance: 0.17 mm), using ZEN imaging software (Carl Zeiss Ltd., UK). All images were analysed with ImageJ 1.53 k software (National Institutes of Health, USA).

### Behavioural analyses

Behavioural tests were performed in the order they are introduced below. Apparatus was cleaned using 70 % ethanol upon completion of each trial, eliminating any residual odour.

Open field test (OFT) was conducted as previously described^10^. Briefly, mice were placed in the centre of the OFT, a grey 50 x 50 x 50 cm apparatus illuminated with low lux (100 lux) lighting. Total travel distance and time spent in the centre of the field was determined at 5 min with a video tracking system (Smart 3.0 tracking software, Panlab, Kent, UK).

Y-maze spontaneous alternation test, a measure of spatial working memory, was performed on the final day of behavioural testing as previously described^10^. Briefly, the Y-maze apparatus comprised white Plexiglas (dimensions 38.5 × 8 × 13 cm, spaced 120° apart) and was illuminated with low lux (100 lux) lighting. Mice were placed in the maze and allowed to explore freely for 7 min while tracking software recorded zone transitioning and locomotor activity (Smart 3.0 tracking software, Panlab, Kent, UK). Spontaneous alternation was calculated using the following formula: Spontaneous Alternation = (Number of alternations/ Total Arm entries - 2) x 100.

The novel object recognition (NOR), a measure of recognition memory, was performed as described previously^10^. Briefly, on day 1 mice were habituated in a grey 50 x 50 x 50 cm apparatus illuminated with low lux (100 lux) lighting, mice were placed into the empty maze and allowed to move freely for 10 min. On day 2, mice were conditioned to a single object for a 10 min period. On day 3, mice were placed into the same experimental area in the presence of two identical objects for 15 min, after which they were returned to their respective cages and an inter-trial interval of 1 h was observed. One familiar object was replaced with a novel object. Mice were placed back within the testing area for a final 10 min. Videos were analysed for a 5 min period, after which if an accumulative total of 15 s with both objects failed to be reached, analysis continued for the full 10 min or until 15 s was achieved. Those not achieving 15 s were excluded from the analysis^73^. A discrimination index (DI) was calculated as follows: DI = (TN−TF)/(TN+TF), where TN is the time spent exploring the novel object and TF is the time spent exploring the familiar object.

### Cell culture

The human cerebromicrovascular endothelial cell line hCMEC/D3 was maintained and treated as described previously^9^. Cells were cultured to confluency in complete endothelial cell growth medium MV2 (PromoCell GmbH, Germany), whereupon VEGF was removed, and cells were further cultured for a minimum of 4 days to enable intercellular tight junction formation prior to experimentation. All cell cultures were used at passages 28-33 to ensure retention of appropriate endothelial characteristics^30^.

### In vitro barrier function assessments

Paracellular permeability and TEER were measured on 100 % confluent hCMEC/D3 cultures polarised by growth on 24-well plate polyethylene terephthalate (PET) transwell inserts (surface area: 0.33 cm^2^, pore size: 0.4 μm; Appleton Woods, UK) previously coated with calf- skin collagen (15 µg/cm^2^ and fibronectin 3 µg/cm^2^; both Sigma-Aldrich, UK). The permeability of hCMEC/D3 cell monolayers to 70 kDa FITC-dextran (2 mg/ml) was measured as described previously^10^. TEER measurements were performed using a Millicell ERS-2 Voltohmmeter (Millipore, Watford, UK) and were expressed as Ω.cm^2^. In all cases, values obtained from cell-free inserts similarly coated with collagen and fibronectin were subtracted from the total values.

### Immunofluorescence analysis

hCMEC/D3 cells were cultured on Lab-Tek™ Permanox™ 8-well chamber slides coated with calf skin collagen (Sigma-Aldrich, UK), prior to immunostaining according to standard protocols^74^ and using a primary antibody directed against zona occludens-1 (ZO-1; 1:100, Thermofisher Scientific, UK) or AF488-conjugated phalloidin (0.165 µM, Thermofisher Scientific, UK). Nuclei were counterstained with DAPI (Sigma-Aldrich, UK). Images were captured using an LSM880 confocal laser scanning microscope (Carl Zeiss Ltd., Cambridge, UK) fitted with 405, 488 and 561 nm lasers and a × 63 oil immersion objective lens (NA, 1.4 mm, working distance, 0.17 mm). Images were captured with ZEN imaging software (Carl Zeiss Ltd., UK) and analysed using ImageJ 1.53k (National Institutes of Health, USA).

### Flow cytometry

Following experimental treatment, hCMEC/D3 cells were detached from growth surfaces as described previously^9^, washed in 0.1 M PBS at 4 °C and fixed by incubation for 15 min at room temperature in 4 % formaldehyde in 0.1 M PBS. For analysis of intracellular phospho-ANXA1, phospho-EGFR or phospho-STAT3 cells were then permeabilised by incubation for 10 min in ice-cold 90 % methanol in 0.1 M PBS. Cells were then incubated at 4 °C for 20 min with 10 % fetal calf serum in 0.1 M PBS, 1 mM CaCl_2_ for 20 min to block non-specific secondary antibody binding; blocking buffer was further supplemented with 0.05 % Triton X-100 to permeabilise cells used for analysis of total EGFR, ANXA1, MMP2 or MMP9. Cells were then incubated with primary antibodies raised against, variously, ANXA1 (rabbit anti-human, 1:1000, ThermoFisher Scientific, UK), phosphotyrosine-21 ANXA1 (rabbit anti-human, 1:1000^72^), EGFR (rabbit anti-human, 1:50, Cell Signalling Technologies, UK), phosphotyrosine-1068 EGFR (mouse anti-human, 1:1000, Cell Signalling Technologies, UK), MMP2 (rabbit anti-human, 1:500, Insight Biotechnology Ltd, UK), MMP9 (mouse anti-human, 1:50, Santa Cruz Biotechnology Inc., USA) or phosphotyrosine-705 STAT3 (mouse anti-human, 1:200, Cell Signalling Technologies, UK) in 1 % FCS in 0.1 M PBS for 20 min on ice. Cells were washed and incubated with either an AF488-conjugated goat anti-rabbit or an AF488-conjugated goat anti-mouse secondary antibody as appropriate (1:300, Thermo-Fisher Scientific, UK) in 1 % FCS, 0.1 M PBS for 20 min on ice. Immunofluorescence was analysed for 10,000 singlet events per treatment using a BD FACSCanto II (BD Biosciences, UK) flow cytometer, with resultant data being analysed with FlowJo 8.0 software (Treestar Inc., CA, USA).

### Gelatinase activity analysis

hCMEC/D3 cells grown on 12-well plates were cultured to confluence in complete endothelial cell growth medium MV2 (PromoCell GmbH, Germany), whereupon serum was removed and cells were subjected to experimental treatment as described. Cell-free medium was collected and concentrated 10-fold by centrifugal filtration (30 kDa cut-off, Pall Corporation, UK) at 14,000 g for 8 min, with resuspension in sterile 0.1 M PBS. Concentrated samples (10 µl/sample) were mixed with 5x non-reducing sample buffer (125 mM Tris.HCl, 4 % sodium dodecyl sulfate (SDS), 20 % glycerol, 0.01 % bromophenol blue, pH 6.8; all Merck, UK) and separated by gel electrophoresis (7.5% acrylamide, 4 mg/ml porcine gelatin; Merck, UK). SDS was removed by washing in 50 mM Tris.HCl, 2.5 % Triton X-100, 5 mM CaCl_2_, 1 µM ZnCl_2_, pH 7.5. Gelatinase activity was determined by incubation for 24 h in 50 mM Tris.HCl, 1 % Triton X-100, 5 mM CaCl_2_, 1 µM ZnCl_2_, pH 7.5 followed by Coomassie staining (0.5 % Coomassie brilliant blue, 40 % methanol, 10 % acetic acid). Gels were visualised using a ChemiDoc MP Imaging System (Bio-Rad Laboratories Ltd., UK). Integrated optical densities were calculated using NIH ImageJ 1.53k (National Institutes of Health, USA).

### EGFR siRNA

hCMEC/D3 cells grown on 24-well plate polyethylene terephthalate (PET) transwell inserts (surface area: 0.33 cm^2^, pore size: 0.4 μm; Appleton Woods, UK) previously coated with calf- skin collagen (15 µg/cm^2^ and fibronectin 3 µg/cm^2^; both Sigma-Aldrich, UK), were transfected with one of three different commercial siRNA sequences designed to target the EGFR or an Allstars negative control siRNA sequence (final concentration 1 nM; all Qiagen Ltd., UK) using INTERFERin transfection reagent according to the manufacturer’s instructions (Polyplus Transfection, France), alongside mock transfected cells. After 96 h, cells were analysed for TEER and paracellular permeability to a 70 kDa FITC-dextran tracer as described above. A proportion of cells were analysed for EGFR expression by flow cytometry as described above (Extended Data Fig. 5).

### ANXA1 shRNA

hCMEC/D3 cells bearing shRNA sequences targeting ANXA1 or a scramble sequence were used as described previously^10^. Prior to use, reduction in ANXA1 expression was confirmed by flow cytometry as described above (Extended Data Fig. 5).

### Cell survival analysis

The potential for pCS-induced cytotoxicity was assessed using the MTT assay. Briefly, cells were treated with pCS (10 µM, 100 µM, 1 mM) for 24 h, prior to administration of MTT (3-(4,5- dimethylthiazol-2-yl)-2,5-diphenyltetrazolium bromide; Merck Life Science UK Ltd., Gillingham, UK) at 500 μg/ml. Cells were incubated at 37 °C for 2 h, medium was removed and resulting crystals were solubilised by incubation for 2 min in dimethyl sulfoxide. Absorbance was read at 540 nm using a CLARIOstar spectrophotometer (BMG Labtech, Ortenberg, Germany), with a reference wavelength at 570 nm.

### LC-MS/MS

Serum samples were diluted with methanol at a ratio of 1:10 (v/v) and placed on dry ice for 10 min. Samples were then centrifuged (5 min, 16,000 *g* at room temp), supernatants filtered using a 0.45 µm PTFE syringe filter and evaporated to dryness using a Savant™ SpeedVac™ High-Capacity Concentrator (Cat. SC210A-230). Dried samples were resuspended in 50 µl of water with 15 µl of L-tryptophan-2,3,3-d_3_ and *p*-toluenesulfonic acid at 50 µg/ml as the internal standards for tryptophan and *p*-cresol metabolites, respectively.

Stock solutions of each metabolite were prepared in methanol (1 mg/ml) and stored at -80 °C. Calibration standards were prepared by pooling all analytes at eight different concentrations and adding the respective internal standards at 50 µg/ml. Calibration standards were run at the beginning, middle and end of each analytical queue.

Two microlitres of the prepared sample was injected and separated using a Waters Acquity UPLC system and Xevo TQ-S Cronos mass spectrometer and ACQUITY UPLC BEH C18 1.7 µm (2.1 x 50 mm) column. The electrospray ionisation operated in positive mode and chromatic separation occurred using a 0.3 ml/min and composition of eluent A (0.1 % formic acid, water) and eluent B (0.1 % formic acid, methanol) at a gradient of 5 % B from 0 to 0.5 min, 5 %-10 % B from 0.5 to 2.5 min, 10 %-15 % B from 2.5-3.5 min, 15 %-35 % B from 3.5 to 4.5 min, 35 % to 45 % B from 4.5 to 6.5 min, 45 % to 55 % B from 6.5 to 7 min, 55 % to 100 % B from 7 to 7.5 min, 100 % B until 10 min, and 5 % B from 10 to 10.1 min to return to initial conditions for equilibration until 14 min. The analyte:internal standard response ratio was used to create a calibration curve and quantify each metabolite.

Sample sizes were calculated to detect differences of 15 % or more with a power of 0.85 and α set at 5 %, calculations being informed by previously published data^10^. Experimental data are expressed as mean ± SEM, with n = 4-9 independent experiments for *in vitro* studies and n=6-10 individual animals for *in vivo*/*ex vivo* studies, see figure legends for details of individual experiments. For *in vivo*/*ex vivo* studies, individual animals were randomly assigned to groups using the =rand() function of Microsoft Excel. Statistical analyses were performed using RStudio 2022.07.1. In all cases, normality of distribution was established using the Shapiro– Wilk test, followed by analysis with two-tailed Student’s *t*-tests to compare two groups or, for multiple comparisons, one- or two-way ANOVA with Tukey’s HSD *post hoc* test as appropriate. A *P* value of <5 % was considered significant.

## Acknowledgements

This work was funded by Alzheimer’s Research UK Pilot Grant No. ARUK-PPG2016B-6. This project has received funding from the European Union’s Horizon 2020 research and innovation programme under grant agreement No 874583. This publication reflects only the authors’ views, and the European Commission is not responsible for any use that may be made of the information it contains. The funders had no role in the study design, data collection or analysis, decision to publish, or preparation of the manuscript.

## Author Contributions

Conceptualisation: LH, SM; Investigation: SNS, TBAK, MGP, EC, GLG, TAJH DV, LH and SM; Resources: DWR, KM, MMY and ES; Writing – original draft: SM; Writing – review & editing: MM, AVS, RCG, DV, LH and SM; Project administration: SM. All authors read and approved the final version of the manuscript.

## Abbreviations

BBB: blood–brain barrier
CKD: chronic kidney disease
EGFR: epidermal growth factor receptor
IS: indoxyl sulfate
MMP: matrix metalloproteinase
pCS: *p*-cresol sulfate
pCG: *p*-cresol glucuronide
SPIA: signalling pathway impact analysis
TEER: transendothelial electrical resistance
ZO-1: zonula occludens-1.

---

**Extended Data Figure 1:**
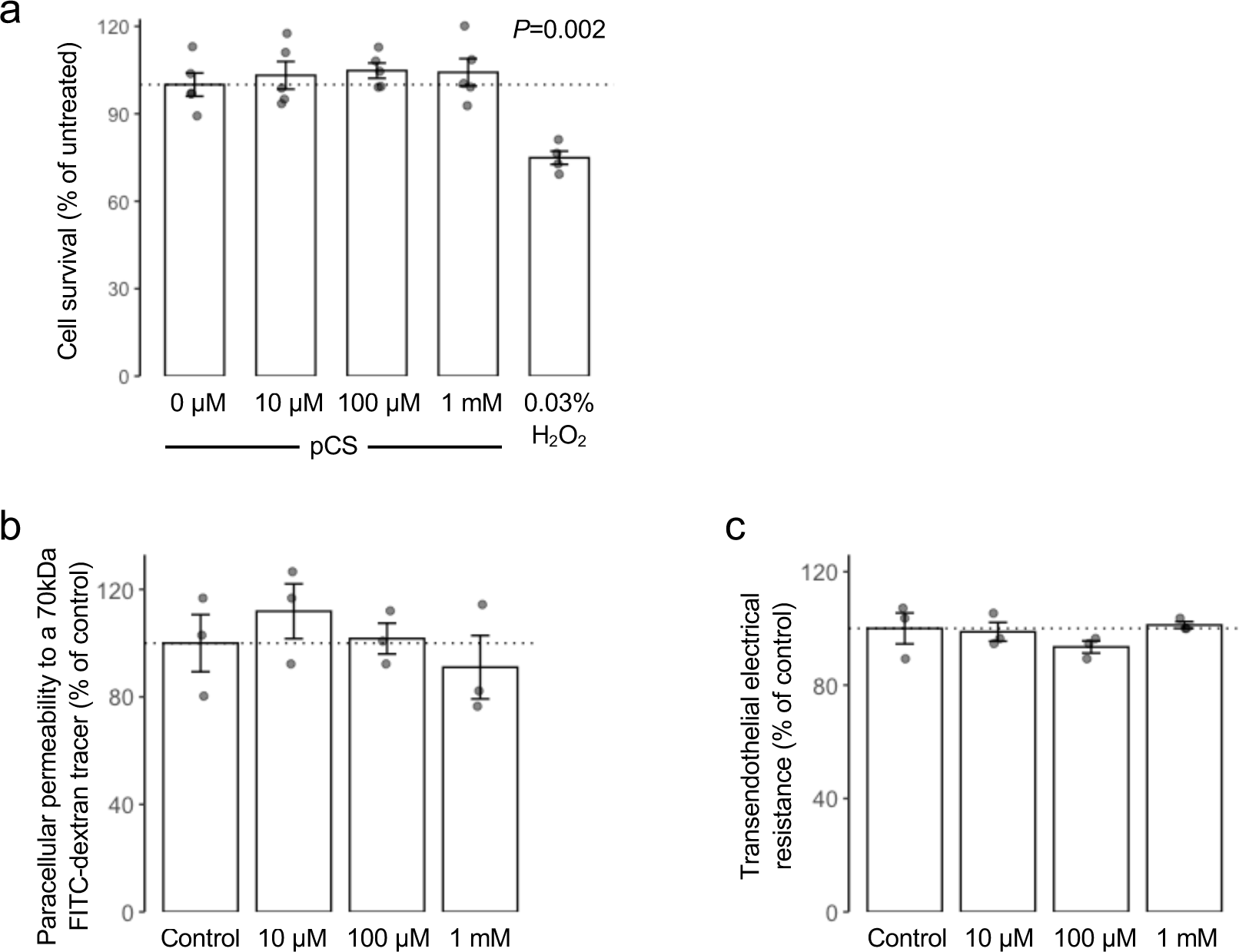
Confirmation of specificity of pCS effects upon endothelial barrier function in hCMEC/D3 cells. a) Treatment of hCMEC/D3 cells with increasing concentrations of pCS (10 µM, 100 µM, 1 mM, 24 h) does not affect cell viability as assessed by MTT assay, in contrast to treatment with 0.03 % H_2_O_2_ (24 h), which caused a significant loss in viability; data are mean ± s.e.m., n = 5 independent experiments. b) Incubation of hCMEC/D3 cell monolayers with KCl (10 µM - 1 mM; 24 h) did not significantly affect paracellular permeability to a 70 kDa FITC-dextran conjugate; data are mean ± s.e.m., n = 3 independent experiments. c) Incubation of hCMEC/D3 cell monolayers with KCl (10 µM – 1 mM; 24 h) did not significantly affect TEER; data are mean ± s.e.m., n = 3 independent experiments.

**Extended Data Figure 2:**
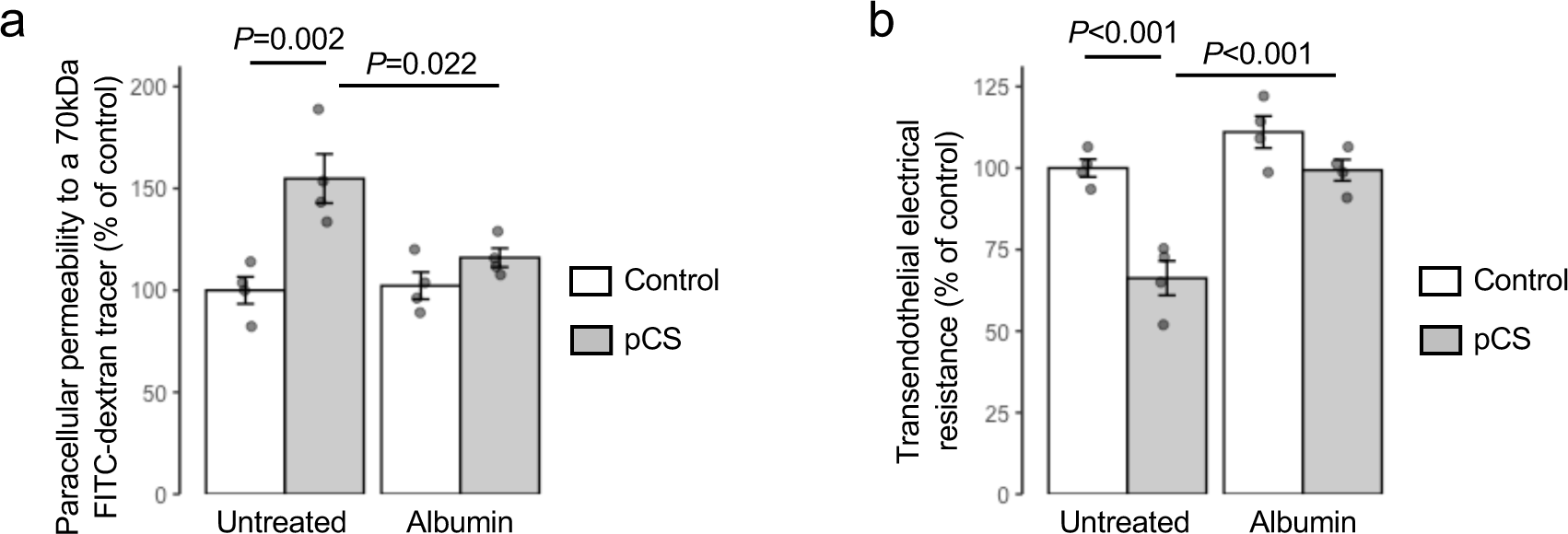
Treatment of hCMEC/D3 cells with albumin prevented the effects of pCS upon endothelial barrier function. a) Incubation of hCMEC/D3 cell monolayers with human albumin (50 mg/ml) in conjunction with pCS (10 µM, 24 h) prevented the effects of pCS upon paracellular permeability to a 70 kDa FITC-dextran conjugate; data are mean ± s.e.m., n = 4 independent experiments. b) Incubation of hCMEC/D3 cell monolayers with human albumin (50 mg/ml) in conjunction with pCS (10 µM, 24 h) prevented the effects of pCS upon trans-endothelial electrical resistance; data are mean ± s.e.m., n = 4 independent experiments.

**Extended Data Figure 3:**
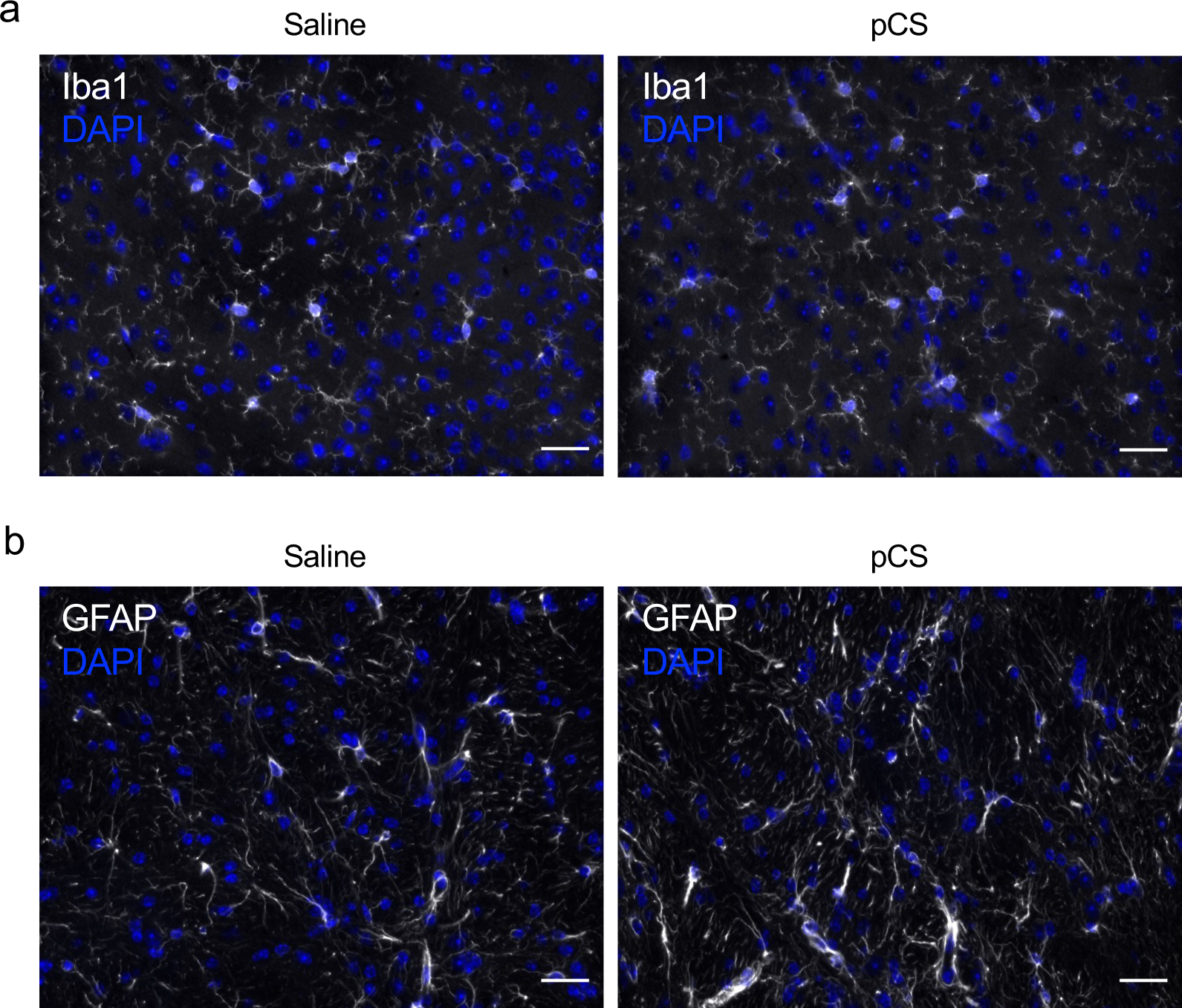
Mice exposed to raised levels of pCS for 4 weeks do not show any obvious changes in cortical microglial or astrocyte morphology or density. Typical immunofluorescent analysis of a) Iba1 expression (microglia) or b) GFAP expression (astrocytes) within the cerebral cortex of male C57Bl/6 mice implanted with minipumps designed to release saline vehicle (n=10) or pCS (n=7) continuously for four weeks. Nuclei are counterstained with DAPI (blue), scale bar = 30 µm.

**Extended Data Figure 4:**
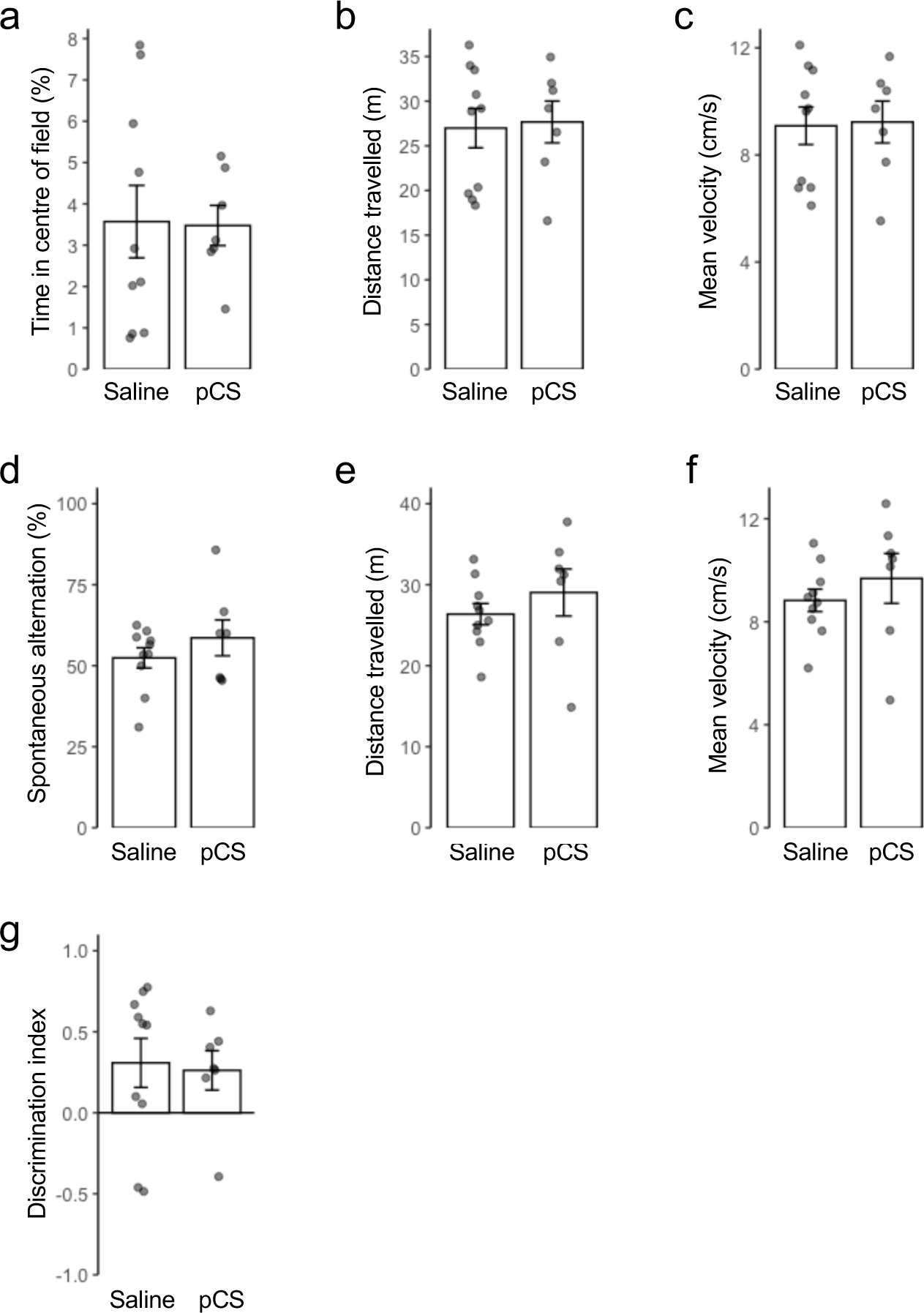
Mice exposed to raised levels of pCS for 4 weeks do not show overt behavioural changes on the open field test, Y-maze or novel object recognition task. Mice implanted with minipumps designed to release saline vehicle or pCS continuously for four weeks were assessed for a) percentage of time in the centre zone, b) distance travelled and c) movement speed in the open field test, d) spontaneous alteration, e) distance travelled and f) mean velocity in the Y-maze task and g) discrimination index in the novel object recognition test. Data are expressed as mean ± s.e.m., *n* = 7-10 mice.

**Extended Data Figure 5:**
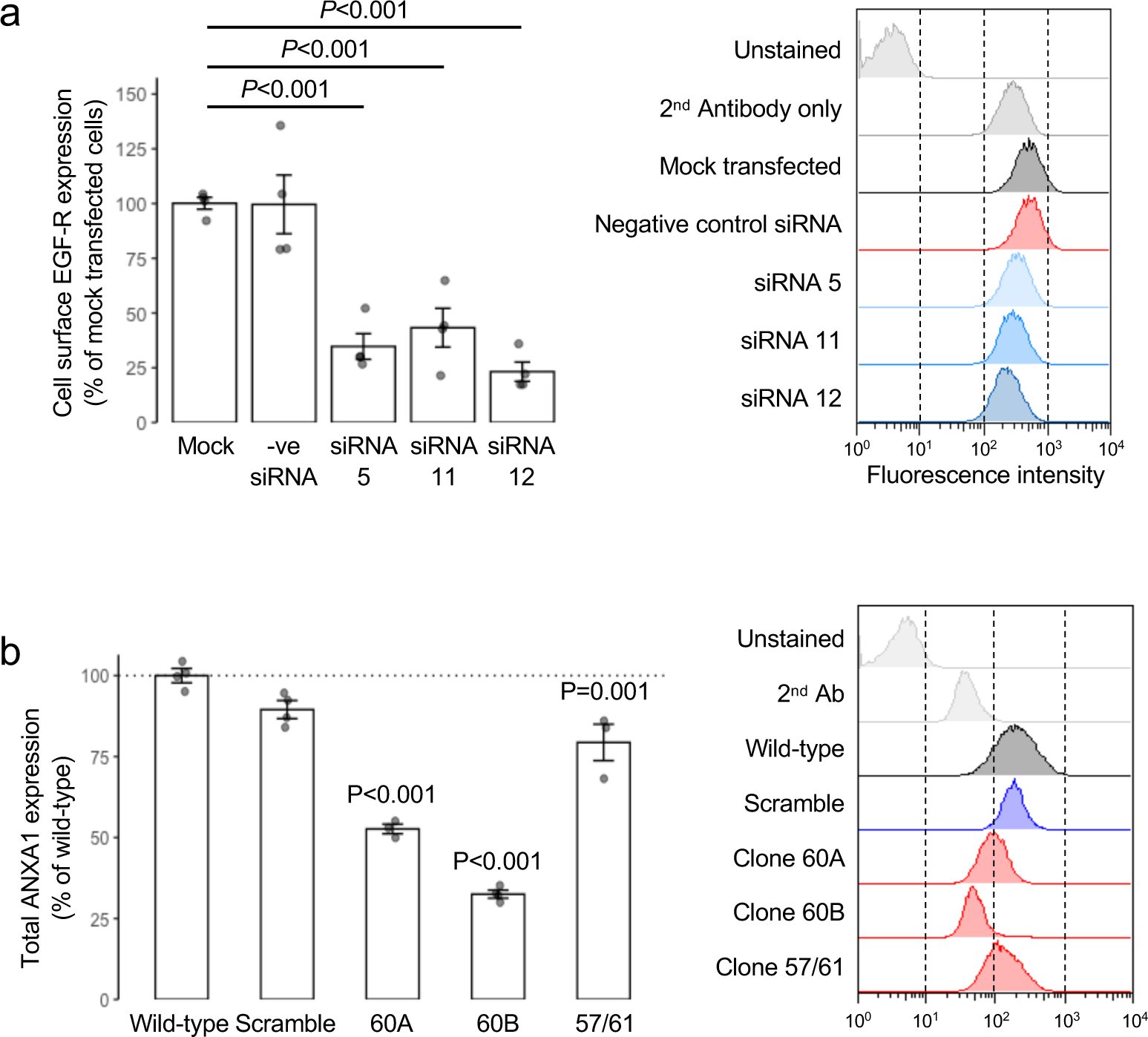
Confirmation of RNA interference knock-down experiments. a) Transfection of hCMEC/D3 cells with siRNA sequences targeting EGFR causes a significant down- regulation of expression 96 h later when compared with mock transfected cells or cells transfected with a non-targeting negative control siRNA sequence; data are mean ± s.e.m., n = 4, representative flow cytometry histograms are shown. b) Stable transfection of hCMEC/D3 cells with shRNA sequences targeting ANXA1 causes a reduction in protein expression, an effect not seen upon transfection with a scramble shRNA sequence; data are expressed as mean ± s.e.m., *n*=4 independent experiments, representative flow cytometry histograms are shown.

**Extended Data Figure 6:**
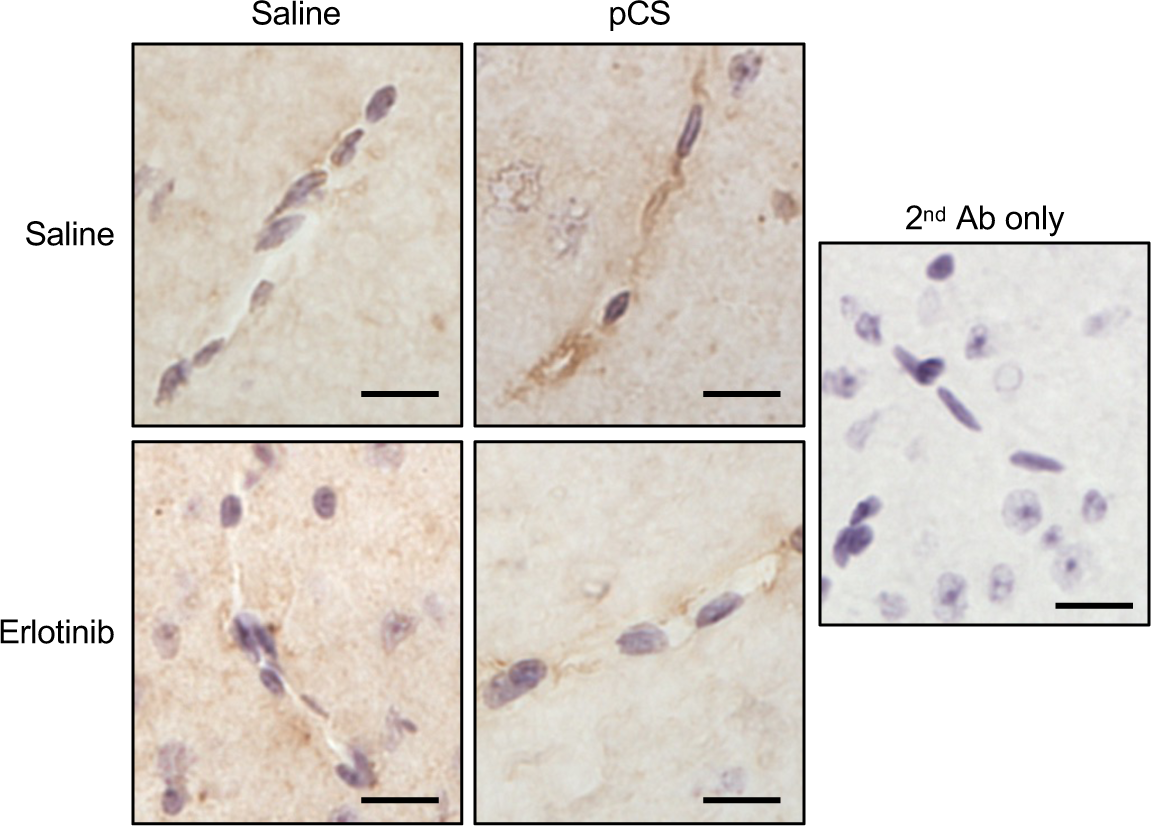
*In vivo* administration of pCS stimulates Tyr21 phosphorylation of ANXA1 in an erlotinib-sensitive manner. Typical immunohistochemical analysis of Tyr21-phosphorylated ANXA1 expression within the cerebral microvasculature of male C57Bl/6 mice treated with pCS (10 mg/kg, i.p. 2 h) with or without erlotinib pre-treatment (50 mg/kg, i.p. 1 h pre-treatment); nuclei are counterstained with haematoxylin, scale bar = 15 µm.

**Extended Data Figure 7:**
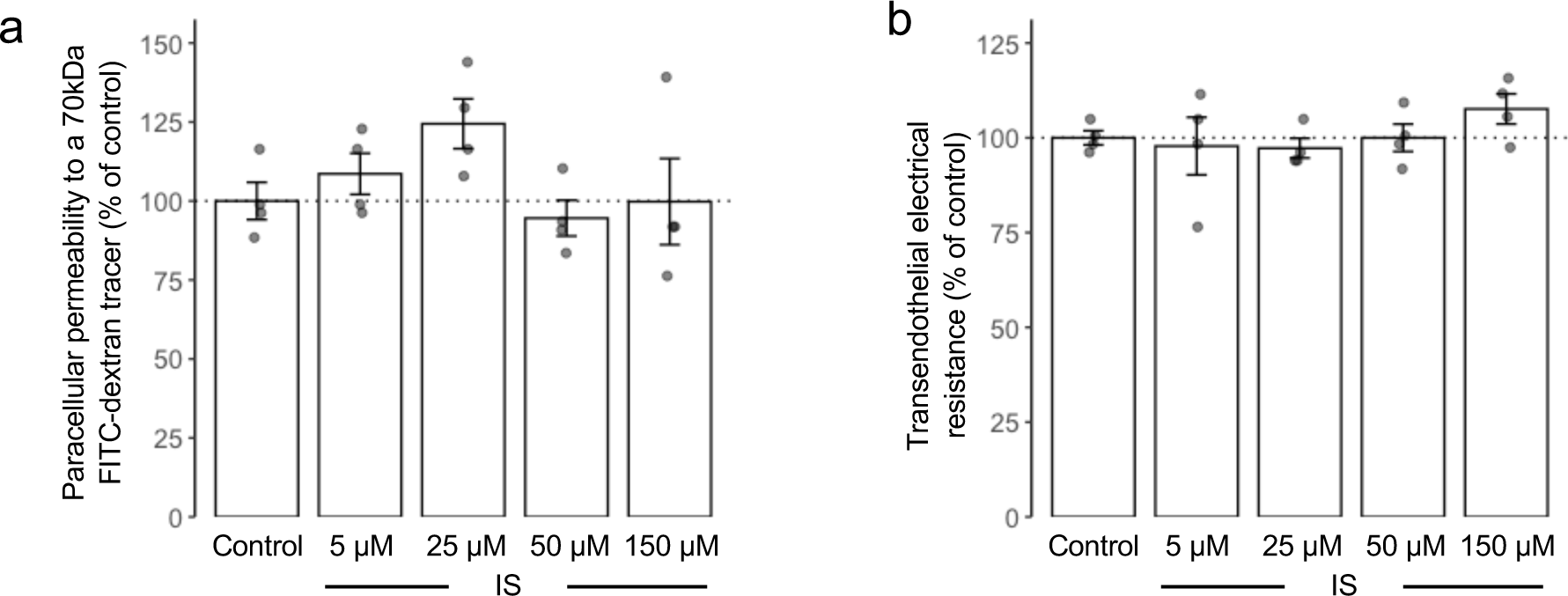
Treatment of hCMEC/D3 cells with indoxyl sulfate (IS) does not affect endothelial barrier functions at physiological or uraemia-associated concentrations. a) Incubation of hCMEC/D3 cell monolayers with IS (5 µM, 25 µM, 50 µM, 150 µM; 24 h) did not significantly affect paracellular permeability to a 70 kDa FITC-dextran conjugate at any concentration tested; data are mean ± s.e.m., n = 4 independent experiments. b) Incubation of hCMEC/D3 cell monolayers with IS (5 µM, 25 µM, 50 µM, 150 µM; 24 h) did not significantly affect TEER at any concentration tested; data are mean ± s.e.m., n = 4 independent experiments.

**Extended Data Table 1:**
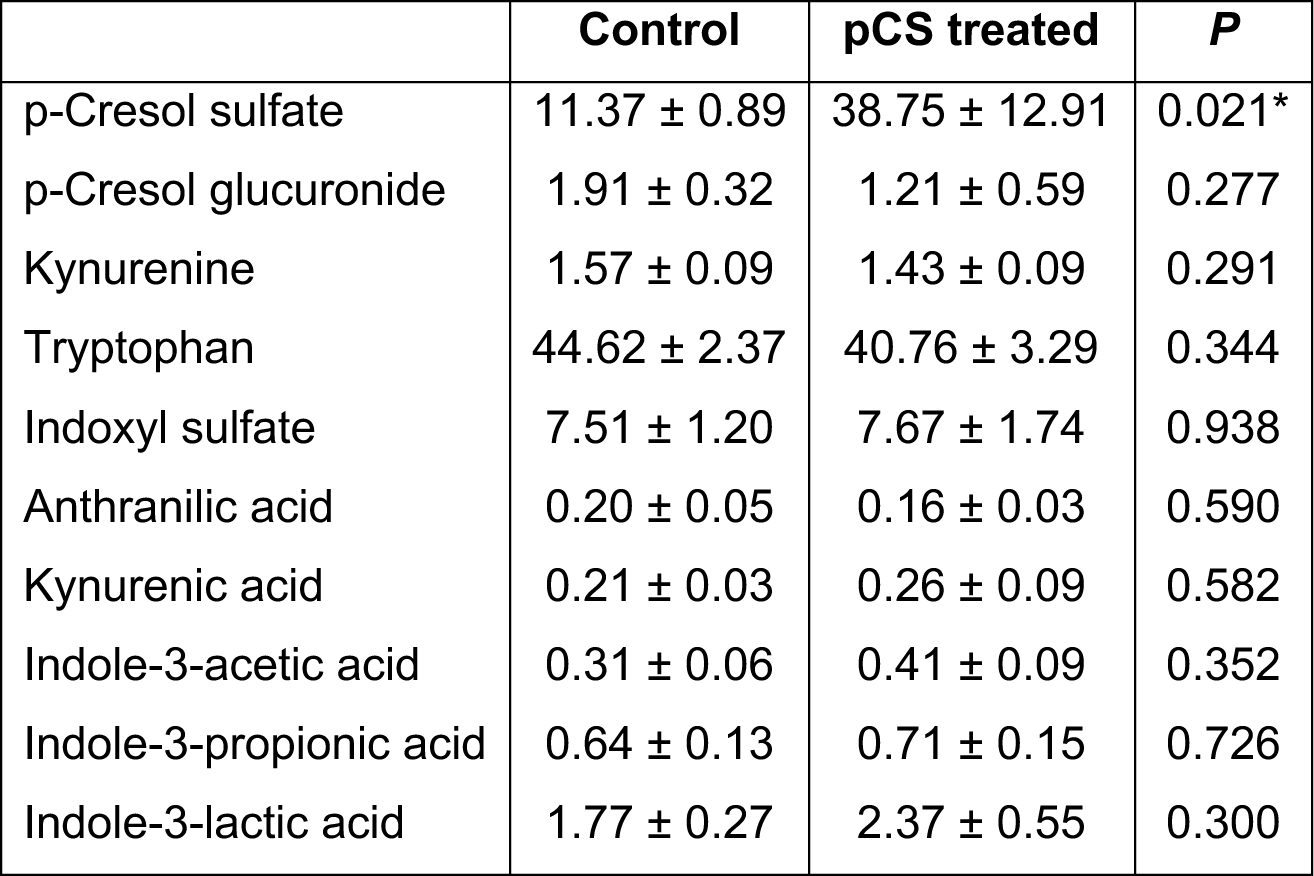
Circulating concentrations (µM) of *p*-cresol and tryptophan metabolites in mice exposed for 8 weeks to pCS or saline *via* osmotic minipumps. Data are mean ± s.e.m., n=7-10, * indicates statistical significance at *P*<0.05 or less

|  | Control | pCS treated | *P* |
| --- | --- | --- | --- |
| p-Cresol sulfate | 11.37 ± 0.89 | 38.75 ± 12.91 | 0.021* |
| p-Cresol glucuronide | 1.91 ± 0.32 | 1.21 ± 0.59 | 0.277 |
| Kynurenine | 1.57 ± 0.09 | 1.43 ± 0.09 | 0.291 |
| Tryptophan | 44.62 ± 2.37 | 40.76 ± 3.29 | 0.344 |
| Indoxyl sulfate | 7.51 ± 1.20 | 7.67 ± 1.74 | 0.938 |
| Anthranilic acid | 0.20 ± 0.05 | 0.16 ± 0.03 | 0.590 |
| Kynurenic acid | 0.21 ± 0.03 | 0.26 ± 0.09 | 0.582 |
| Indole-3-acetic acid | 0.31 ± 0.06 | 0.41 ± 0.09 | 0.352 |
| Indole-3-propionic acid | 0.64 ± 0.13 | 0.71 ± 0.15 | 0.726 |
| Indole-3-lactic acid | 1.77 ± 0.27 | 2.37 ± 0.55 | 0.300 |

**Extended Data Table 2:** Outputs from DESeq2 analysis of differentially expressed genes.

**Extended Data Table 3:** Summary network statistics for transcriptomic data mapped to KEGG pathways.

